# Targeted dephosphorylation of Tau at the endogenous level promotes its association with microtubules

**DOI:** 10.64898/2026.09.07.749800

**Authors:** Gajanan Sathe, Francesco Berterame, Tyrell Cartwright, Ria Spallek, Jin-Feng Zhao, Ayush R. Bagale, Robert Gourlay, Nicola T. Wood, Thomas J. Macartney, Gopal P. Sapkota

## Abstract

Hyperphosphorylation of Tau is a contributory factor for Tauopathies, which constitute a spectrum of neurodegenerative disorders including Alzheimer’s disease (AD). Inhibitors of Tau kinases or activators of the Tau phosphatase PP2A to reverse Tau hyperphosphorylation have shown limited therapeutic promise, due primarily to a lack of Tau selectivity. An innovative strategy to potentially circumvent these limitations would be to selectively target the dephosphorylation of phospho-residues on Tau. In this study, we provide evidence for targeted dephosphorylation of phospho-Tau at the endogenous level, including those known to promote Tau aggregation. By recruiting PPP1CA and PPP2CA Affinity-directed (Ad)-Phosphatases to Tau, we demonstrate highly selective targeted dephosphorylation of multiple phospho-residues on Tau in SK-N-MC neuroblastoma cells, leading to its stabilization in microtubules. Moreover, by using a heterobifunctional molecule termed BDPIC (bromoTAG-dTAG proximity-inducing chimera) in cells harbouring bromoTAG and dTAG knockins on Tau and PPP2CA, respectively, we demonstrate highly selective dephosphorylation phospho-residues on Tau.

**Summary:** By employing induced-proximity approaches, we recruit phosphatases to endogenous Tau for targeted dephosphorylation.

## Introduction

Neurofibrillary tangles (NFTs) composed of hyperphosphorylated Tau protein, which is encoded by the microtubule-associated protein tau (MAPT) gene, are characteristic features of a group of neurodegenerative diseases called tauopathies, which includes AD ^1–3^. Aberrantly enhanced phosphorylation of over 40 serine, threonine or tyrosine residues on Tau by different protein kinases has been reported from AD and other tauopathy brain samples compared to controls ^4,5^. In normal physiology, Tau is a soluble microtubule-associated protein that promotes assembly and stability of microtubules (MT) ^6^. Hyperphosphorylation of Tau causes a reduction in its ability to bind to microtubules ^7^. Indeed, disruption of the MT network is frequently observed in tauopathies, and this is thought to result from Tau loss-of-function due to its hyperphosphorylation ^8^. Progressive Tau hyperphosphorylation and aggregation in patient brains are key hallmarks of AD ^9,10^. Many protein kinases, including glycogen synthase kinase-3β (GSK-3β), cyclin-dependent kinase-5 (CDK-5), extracellular signal regulated kinase 1/2 (ERK1/2), p38 mitogen activated protein kinase α (p38α) and casein kinase 1 (CK1) are reported to phosphorylate Tau ^11^. Similarly, protein phosphatase 2A (PP2A) is known to reverse Tau phosphorylation ^12–17^. In normal physiology, an intricate balance in activities between the Tau kinases and phosphatases controls homeostasis, while amplifications of some Tau kinases or dampening of the PP2A activity has been reported in AD brains ^18–20^.

Although the precise causal relationship between Tau hyperphosphorylation and AD pathology remains to be fully defined, hyperphosphorylated Tau has been reported to be toxic to neurons, thereby promoting neuronal death ^21^. Hyperphosphorylation of Tau is not only associated with increased propensity for Tau to aggregate ^22–24^ but also shown to directly contribute to progressive AD pathology ^25,26^. Alternative splicing results in expression of multiple Tau variants ^27^, however in this manuscript the residue numbers refer to the human 2N4R variant of Tau. Phosphorylation of some key sites, including Y18, Thr175, Thr181, Ser199, Ser202, Thr205, Thr208, Thr212, Ser396, Ser404, and Ser422, or mimicking phosphorylation at these sites has been reported to promote Tau aggregation ^23,28–35^. Paired helical filaments (PHFs), the main structural units of NFTs, feature Tau hyperphosphorylated at residues, such as Ser396 and Ser404, which serve as faithful diagnostic and mechanistic biomarkers of AD ^36^. GSK3β-mediated phosphorylation of the PHF-1 epitope (Ser396/Ser404) was reported to drive Tau aggregation into Alzheimer-like filaments ^37^. Preclinical studies targeting PHF-1 with systemic anti-PHF-1 antibodies, single-chain variable fragments, or adenoviral-delivered intrabodies have demonstrated reduced Tau pathology and improved phenotypes in Tau transgenic mice ^38–40^. In MAPT-P301S mice, hippocampal PHF-1 expression via adenoviral delivery prevented atrophy and Tau accumulation ^41^. Other phospho-sites, such as pSer202 and pThr205, are common in PHFs and are used to stage AD pathology ^42^. A recent study reported that phosphorylation of a few residues on Tau, including Thr205 and Thr181, was critical for triggering the hyperphosphorylation of Tau and mutating these ‘master sites’ not only prevented Tau hyperphosphorylation but slowed cognitive impairment in AD rodents ^35^. Indeed, pThr181 on Tau has been used as a biomarker for the efficacy of LEQEMBI, one of just two drugs recently approved for AD therapy ^43^. Phosphorylation at Y18 ^44^, S199 ^45^, S208 ^23^, T212 ^46^, and S422 ^47^ has been associated with AD. Key kinases reported to be responsible for phosphorylating Tau include GSK3β, CDK5, PKA, CaMKII, Fyn, BRSK, MARK, MAPK, LRRK2 and CK1 ^44,48,49^, although precisely how they regulate the phosphorylation of normal and pathogenic Tau remain to be elucidated fully (Fig. S1). With overwhelming evidence linking Tau hyperphosphorylation to AD pathology, reducing the levels of hyperphosphorylated Tau in neurons has been a key therapeutic intervention strategy against AD and other tauopathies. Indeed, several kinase inhibitors, including those targeting GSK3β, p38α and ERK1/2, have undergone AD clinical trials but have not succeeded primarily due to adverse effects, as these kinases are known to regulate many other physiological processes ^11^. Similarly, activating the phosphatase PP2A with tricyclic sulfonamides was shown to reduce Tau hyperphosphorylation and ameliorate AD pathogenesis ^50^, although activating PP2A would be expected to potentially also dephosphorylate over 2000 PP2A substrates in cells ^51^ and could therefore cause many adverse effects. An alternative approach to potentially mitigating the adverse effects of kinase inhibitors and phosphatase activators would be to redirect protein phosphatases directly to Tau through proximity induction to effectuate targeted dephosphorylation of hyperphosphorylated Tau.

Previously, we reported the development of the AdPhosphatase (Affinity-directed Phosphatase) system for highly specific targeted dephosphorylation of phospho-proteins of interest, including FAM83D and ULK1, at the endogenous level ^52^. Recently, we described a small molecule-based approach for targeted dephosphorylation of phospho-proteins, including SMAD3 and TFEB, at the endogenous level ^53,54^. Here, we describe the development and characterization of the AdPhosphatase system for the targeted dephosphorylation of Tau at the endogenous level. Additionally, we also evaluate proof-of-concept for targeted dephosphorylation of Tau at the endogenous level by using the heterobifunctional molecule bromoTAG-dTAG proximity-inducing chimera (BDPIC) in cells harbouring knockins of dTAG and bromoTAG on *MAPT* and *PPP2CA* loci, respectively.

## Results

### Generation of a Tau-GFP knock-in (KI) human neuroblastoma SK-N-MC cell line

Progressive Tau hyperphosphorylation leading to its aggregation into NFTs are key hallmarks of AD (Fig. S1). To choose a cell line model to investigate targeted dephosphorylation of Tau at the endogenous level, we first screened a panel of routinely cultured human cell lines for Tau protein expression by immunoblotting (Fig. 1A). We also interrogated the DepMap proteomics database covering 379 cell lines from the Cancer Cell Line Encyclopedia (CCLE) ^55^ for relative Tau and PPP2CA protein abundance (Fig. 1B, S2). Notably, our immunoblotting data revealed a remarkable concordance with the DepMap data on relative Tau (gene *MAPT*) protein abundance in different cell lines (Fig. 1A&B). Elevated expression of multiple Tau isoforms, including big Tau ^27,56^, was observed in SK-N-MC glioblastoma cells in comparison to other cell lines tested (Fig. 1A). SK-N-MC cells were also identified among those cells exhibiting the highest relative abundance of PPP2CA protein (Fig. S2). Therefore, we chose SK-N-MC cells for further experiments. To facilitate the study of Tau dephosphorylation using the AdPhosphatase system ^52^, we employed CRISPR/Cas9 technology to generate a homozygous knock-in of green fluorescent protein (GFP) cDNA at the C-terminus of the native *MAPT* gene. Many isoforms of Tau are evident in adult human cells due to alternative splicing, but all isoforms share the identical C-terminus ^27^. The homozygous insertion of GFP at the native *MAPT* locus was confirmed by genomic sequencing (Fig. S3). The homozygous GFP knock-in on Tau was further confirmed by immunoblotting using TAU40 antibody, where the appearance of higher-molecular-weight Tau-GFP species in Tau-GFP knock-in (KI) SK-N-MC cells was evident, with concurrent disappearance of native-molecular-weight Tau species seen in wild-type (WT) SK-N-MC cells (Fig. 1C). Furthermore, anti-GFP immunoprecipitation (IP) robustly co-precipitated all isoforms of Tau-GFP from KI but not from WT SK-N-MC cells, and accordingly Tau was completely immune-depleted from post-IP flow-through extracts from KI cells but not WT cells compared to pre-IP input extracts (Fig. 1C). We analysed the levels of Tau phosphorylation on different residues in WT and KI SK-N-MC cells. Notably, we observed no significant alterations in the basal phosphorylation levels at S396 and S202/T205 on Tau between WT and KI cells. However, a modest elevation in levels of phosphorylation at S404 and T181 on Tau was detected in KI compared to the WT cells (Fig. 1D). The KI cells are derived from a single cell clone, whereas the WT cells represent a heterogeneous cell population. Therefore, the slight increase in pS404 and pT181 observed in KI SK-N-MC compared to WT cells is likely due to the clonal variation.

**Figure 1:**
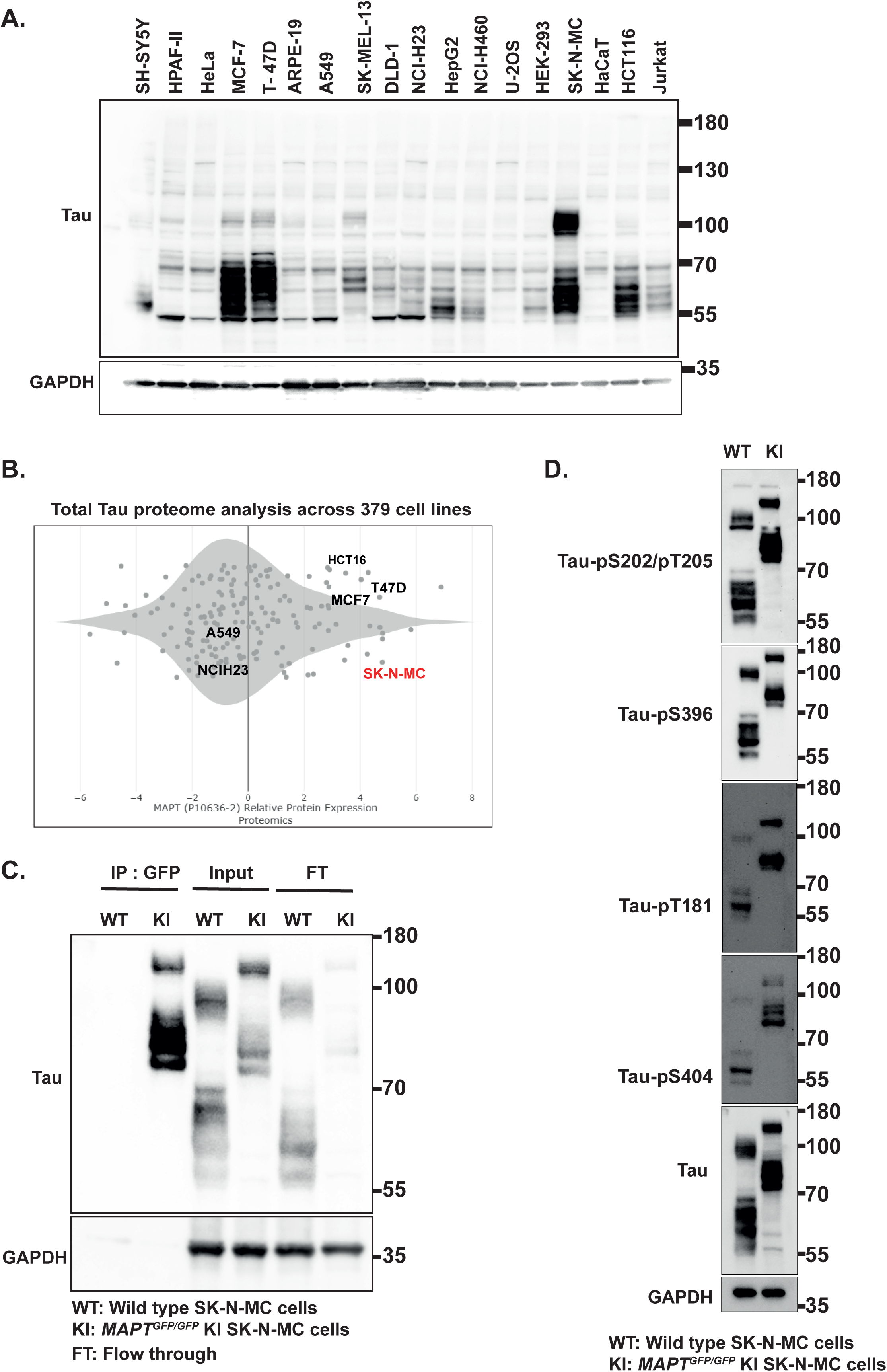
Generation of Tau-GFP knockin (KI) SK-N-MC cells by CRISPR/Cas9. **(A)** Panel of the indicated cell lines screened for the expression of Tau protein using immunoblotting. Extracts (30 μg protein) were resolved on the 4-12% gradient SDS-PAGE, transferred to nitrocellulose membranes and subjected to immunoblotting analysis using the indicated antibodies. **(B)** Cancer Cell Line Encyclopedia (CCLE) proteomics data were mined using DepMap portal and the relative abundance of Tau across all cancer cells were plotted. **(C)** Wild type (WT) or *MAPT^GFP/GFP^* SK-N-MC cells were lysed, and extracts (1 mg protein) were subjected to immunoprecipitation (IP) with GFP-trap beads. Pre-IP input or post-IP flowthrough (FT) extracts (30 μg protein) or GFP IPs were resolved on the 4-12% gradient SDS-PAGE, transferred to nitrocellulose membranes and subjected to immunoblotting analysis using the indicated antibodies. **(D)** Wild type (WT) or *MAPT^GFP/GFP^* SK-N-MC cells were lysed, and extracts (30 μg protein) were resolved on the 4-12% gradient SDS-PAGE, transferred to nitrocellulose membranes and subjected to immunoblotting analysis using the indicated antibodies. All blots are representative of at least 3 independent biological replicates.

### Targeted dephosphorylation of Tau-GFP by the aGFP_6M_-PPP2CA AdPhosphatase

Next, we sought to test whether the aGFP_6M_-PPP2CA Affinity-Directed Phosphatase (AdPhosphatase) ^52^ was capable of targeted dephosphorylation of Tau-GFP in SK-N-MC KI cells. The AdPhosphatase system is a targeted protein dephosphorylation platform that redirects phosphatase activity toward a specific protein of interest ^52^. It comprises a phosphatase catalytic subunit (e.g. PPP1CA or PPP2CA) fused to an antigen-stabilized anti-GFP nanobody (aGFP_6M_), which selectively binds GFP-tagged target proteins. Recruitment of the phosphatase to the target protein then promotes targeted dephosphorylation. The aGFP_6M_ nanobody, which is fused to PPP2CA, was engineered in such a way that it is rapidly degraded when it is not bound to GFP but becomes stabilized upon binding to GFP or GFP-tagged target proteins ^57^. Consequently, the aGFP_6M_-PPP2CA expression levels are expected to closely track the abundance of the target Tau-GFP protein in cells, thereby limiting the accumulation of excess aGFP_6M_-PPP2CA phosphatase within cells. This design offers a key advantage by reducing the potential off target dephosphorylation events in cells that may otherwise arise from phosphatase overexpression (Fig. 2A). Cells were transduced with retroviruses encoding FLAG-aGFP_6M_ control, FLAG-aGFP_6M_-PPP2CA, catalytically inactive FLAG-aGFP_6M_-PPP2CA^H118Q^ mutant, or FLAG-PPP2CA control (Fig. 2B-D). First, in anti-GFP IPs from cell extracts that efficiently precipitated Tau-GFP, robust co-precipitation of FLAG-aGFP_6M_, FLAG-aGFP_6M_-PPP2CA and FLAG-aGFP_6M_-PPP2CA^H118Q^ but not FLAG-PPP2CA control was observed, with almost complete depletion observed in post-IP flow-through extracts (Fig. 2B). Reciprocally, anti-FLAG IPs from extracts that efficiently precipitated FLAG-aGFP_6M_, FLAG-aGFP_6M_-PPP2CA, FLAG-aGFP_6M_-PPP2CA^H118Q^ also precipitated Tau-GFP but anti-FLAG IPs from untransduced or FLAG-PPP2CA transduced control cells did not co-precipitate Tau-GFP (Fig. 2C). Collectively, the data confirm the interaction between aGFP_6M_-AdPhosphatases and Tau-GFP in cells. Crucially, the data also establishes that there is no interaction between Tau-GFP and FLAG-PPP2CA in the absence of aGFP_6M_. Next, we conducted immunoblotting to examine the levels of phosphorylation of some known phospho-sites on Tau, including S396, S404, S202/T205 and T181 relative to total Tau levels, as progressive hyperphosphorylation at these sites has been implicated in the pathogenesis of AD ^25,36,58^. The levels of phosphorylation at all these sites were substantially reduced in cells transduced with FLAG-aGFP_6M_-PPP2CA compared to un-transduced controls or those transduced with FLAG-aGFP_6M_, FLAG-aGFP_6M_-PPP2CA^H118Q^ and FLAG-PPP2CA controls. To assess the specificity of the AdPhosphatase approach, we also monitored the phosphorylation status of AKT at Ser473, a reported PP2A-regulated phosphorylation site ^59,60^. No detectable changes in abundance of AKT pSer473 was observed under any of the conditions tested, indicating that the AdPhosphatase system does not appear to affect native PP2A substrates (Fig. 2D). These data demonstrate targeted dephosphorylation of Tau-GFP by the active aGFP_6M_-PPP2CA AdPhosphatase. To definitively demonstrate the targeted nature of dephosphorylation of Tau-GFP by FLAG-aGFP_6M_-PPP2CA catalytic activity, we immunoprecipitated the AdPhosphatases or the controls using anti-FLAG antibody and examined the phospho-residues on co-precipitating Tau-GFP (Fig. 2E). Indeed, the levels of all phospho-sites on Tau-GFP co-precipitating with Flag-aGFP_6M_-PPP2CA were almost completely abolished compared to those observed on Tau-GFP co-precipitating with FLAG-aGFP_6M_ or FLAG-aGFP_6M_-PPP2CA^H118Q^ (Fig. 2E). As expected, no Tau-GFP co-precipitated with anti-FLAG IPs from untransduced cell extracts or those transduced with FLAG-PPP2CA (Fig. 2E). Our findings underscore the effectiveness of the Flag-aGFP_6M_-PPP2CA Adphosphatase system in achieving targeted dephosphorylation of Tau-GFP at the endogenous level.

**Figure 2:**
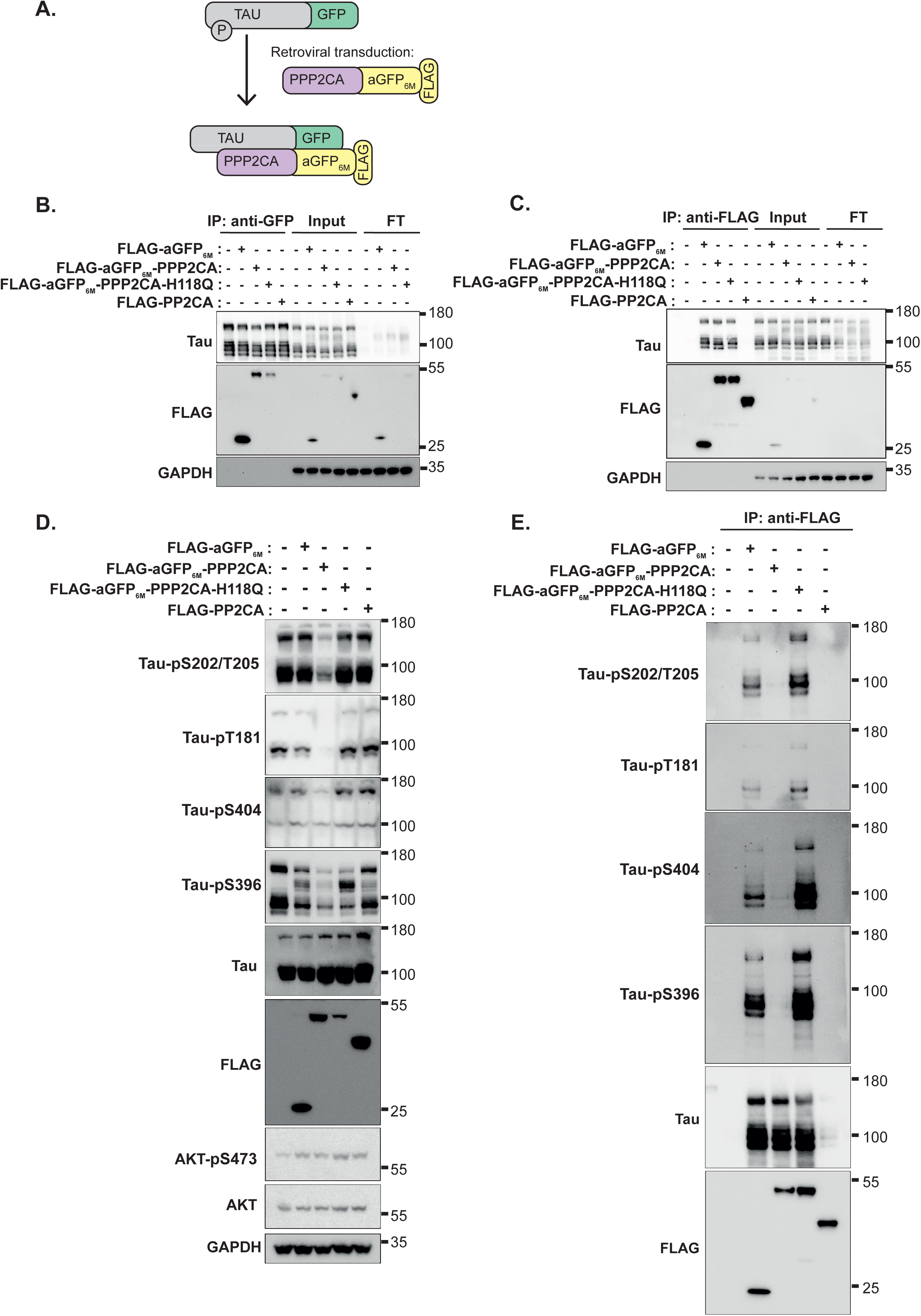
aGFP_6M_-PPP2CA AdPhosphatase dephosphorylates phospho-Tau-GFP. **(A)** Schematic representation of anti-GFP nanobody (aGFP_6M_)-directed recruitment of PPP2CA to C-terminally GFP-tagged Tau to mediate Tau-GFP dephosphorylation. **(B)** Control *MAPT^GFP/GFP^* KI SK-N-MC cells or those stably expressing FLAG-aGFP_6M_, FLAG-aGFP_6M_-PPP2CA, FLAG-aGFP_6M_-PPP2CA^H118Q^, or FLAG-PPP2CA were lysed, and extracts (200 μg protein) subjected to anti-GFP IP. Pre-IP input or post-IP flowthrough (FT) extracts (20 μg protein) or GFP IPs were resolved on the 4-12% gradient SDS-PAGE, transferred to nitrocellulose membranes and subjected to immunoblotting analysis using the indicated antibodies. **(C)** As in (B) except the extracts were subjected to anti-FLAG IPs. **(D)** Control *MAPT^GFP/GFP^* KI SK-N-MC cells or those stably expressing FLAG-aGFP_6M_, FLAG-aGFP_6M_-PPP2CA, FLAG-aGFP_6M_-PPP2CA^H118Q^, or FLAG-PPP2CA were lysed, and extracts (20 μg protein) were resolved on the 4-12% gradient SDS-PAGE, transferred to nitrocellulose membranes and subjected to immunoblotting analysis using the indicated antibodies. **(E)** Extracts (200 μg protein) from (D) were subjected to anti-FLAG IPs, which were resolved on the 4-12% gradient SDS-PAGE, transferred to nitrocellulose membranes and subjected to immunoblotting analysis using the indicated antibodies. All blots are representative of at least 3 independent biological replicates.

### Unbiased global phospho-proteomic analysis demonstrates specificity of the FLAG-aGFP_6M_-PPP2CA AdPhosphatase system

To assess the specificity of FLAG-aGFP_6M_-PPP2CA AdPhosphatase-mediated dephosphorylation of Tau-GFP, we undertook an unbiased quantitative phospho- and total-proteomic analyses in SK-N-MC Tau-GFP KI cells transduced with either FLAG-aGFP_6M_-PPP2CA or the catalytically inactive FLAG-aGFP_6M_-PPP2CA^H118Q^ mutant (Fig. S4A). Total proteome analysis identified a total of 8182 proteins, of which only 16 proteins exhibited a significant change in abundance exceeding a 2-fold threshold between cells expressing FLAG-aGFP_6M_-PPP2CA versus those expressing the FLAG-aGFP_6M_-PPP2CA^H118Q^ mutant (Fig. 3A, Table 1), suggesting that the expression of the two AdPhosphatases does not induce substantial alterations in total protein abundance. Phospho-peptide analysis following enrichment from the same samples identified 36,823 unique phospho-peptides, corresponding to 5732 proteins (Fig. 3B, Tables 2). Among these, the levels of phospho-peptides corresponding to 21 proteins were significantly downregulated by >2-fold, while six were upregulated by >2-fold in Tau-GFP KI cells expressing FLAG-aGFP_6M_-PPP2CA compared to those expressing FLAG-aGFP_6M_-PPP2CA^H118Q^. Notably, within the significantly downregulated phospho-peptides in FLAG-aGFP_6M_-PPP2CA expressing cells, five were on Tau, including those harbouring pS493 and pS462 (Fig. 3B, Table 2). Beyond Tau, Gene Ontology ^61^ and KEGG pathway ^62^ analyses revealed that several other identified phospho-peptides correspond to proteins involved in RNA processing and splicing, including DDX23 ^63^, USP39 ^64^, IWS1 ^65^, THOC5 ^66^, COIL^67^, SCAF1 ^68^, PRPF38B ^69^, HTATSF1 ^70^, ACIN1 ^71^, BCLAF1 ^72^, and ZC3H13 ^73^, and network analysis of these proteins shows their involvement in RNA splicing, mRNA processing, and RNA binding (Fig. S4B, S5). The downregulation of these RNA processing and splicing-related phospho-peptides in cells expressing FLAG-aGFP_6M_-PPP2CA suggests a potential role for the AdPhosphatase system in modulating proteins involved in the regulation RNA processing and splicing (Fig S4B, S5).

**Figure 3:**
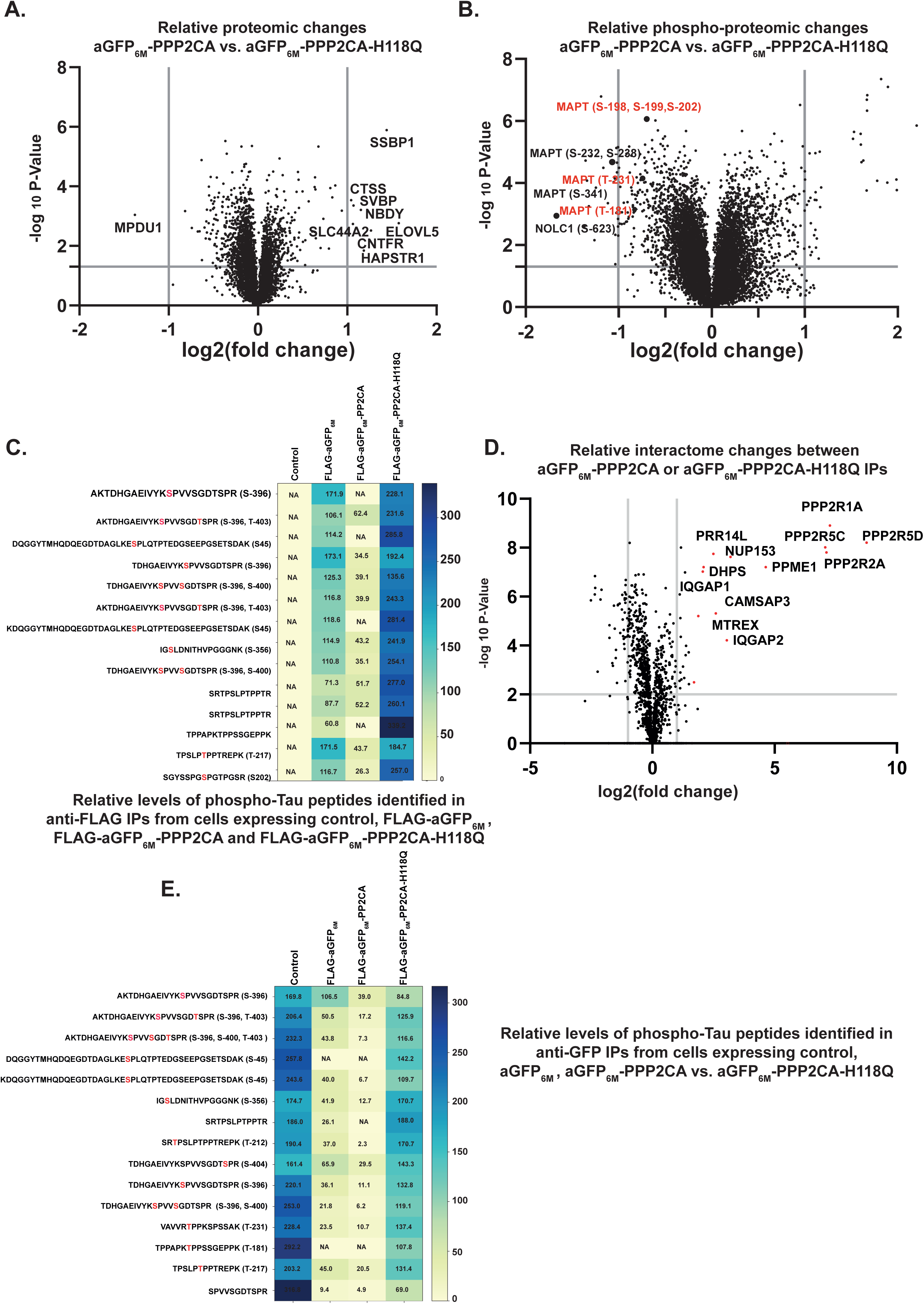
Analysis of the proteomic and phospho-proteomic changes caused by the aGFP_6M_-AdPhosphatase system. **(A)** TMT-labelled quantitative global proteomic analysis of *MAPT^GFP/GFP^*KI SK-N-MC cells expressing FLAG-aGFP_6M_-PPP2CA vs. the FLAG-aGFP_6M_-PPP2CA-H118Q mutant AdPhosphatases. The X-axis represents log2(fold change) in abundance and Y-axis represents -log10(P-Value). **(B)** TMT-labelled quantitative global phospho-proteomic analysis of *MAPT^GFP/GFP^* KI SK-N-MC cells expressing FLAG-aGFP_6M_-PPP2CA vs. the FLAG-aGFP_6M_-PPP2CA^H118Q^ mutant AdPhosphatases. The X-axis represents log2(fold change) in abundance and Y-axis represents -log10(P-Value). The Tau phospho-sites that were significantly downregulated by FLAG-aGFP_6M_-PPP2CA are indicated. **(C)** *MAPT^GFP/GFP^* KI SK-N-MC cells or those stably expressing FLAG-aGFP_6M_, FLAG-aGFP_6M_-PPP2CA or FLAG-aGFP_6M_-PPP2CA^H118Q^ AdPhosphatases were lysed, and extracts subjected to anti-FLAG IP. The IPs were subjected to trypsin digestion and analysed on mass spectrometry for phospho-peptide analysis using Data Dependent Analysis (DDA). The heatmap represents the relative changes in abundance of the specific phospho-Tau peptides identified. The precise phospho-site identified for each peptide is annotated. Where it is not annotated, one of multiple residues could be phosphorylated. **(D)** As in (C) except that FLAG IPs were performed in *MAPT^GFP/GFP^* KI SK-N-MC cells stably expressing FLAG-aGFP_6M_-PPP2CA or FLAG-aGFP_6M_-PPP2CA^H118Q^ AdPhosphatases and subjected to trypsin digestion and analysed on mass spectrometry for proteome analysis using Data Independent Analysis (DIA). The X-axis represent the log2(fold change) in abundance of identified proteins and Y-axis represents -log10(P-Value). **(E)** As in (C) except that GFP-trap beads were used to IP Tau-GFP from cells and the GFP IPs were subjected to trypsin digestion and analysed on mass spectrometry for phospho-Tau peptide analysis using Data Dependent Analysis (DDA).

### Mapping Tau-GFP phospho-sites targeted for dephosphorylation by the FLAG-aGFP_6M_-PPP2CA AdPhosphatase

Without enrichment of Tau-GFP, the global phospho-proteomic analysis above identified only a few of the reported Tau phospho-sites. To identify the full portfolio of AdPhosphatase-mediated dephosphorylation residues on Tau-GFP, we enriched Tau-GFP that was associated with FLAG-aGFP_6M_ control, FLAG-aGFP_6M_-PPP2CA or the FLAG-aGFP_6M_-PPP2CA^H118Q^ mutant via anti-FLAG IPs (Fig. 1E) and performed phosphoproteomic analysis on the co-precipitated Tau-GFP (Fig. 3C). In total, phosphorylation of 17 sites on Tau-GFP that were identified in anti-FLAG IPs from aGFP_6M_ and the catalytically inactive FLAG-aGFP_6M_-PPP2CA^H118Q^ mutant controls were absent or lower in abundance in IPs from FLAG-aGFP_6M_-PPP2CA (Fig. 3C, Table 3), indicating the extent of targeted dephosphorylation of Tau with PPP2CA. As expected, no Tau-GFP or phospho-Tau peptides were evident in anti-FLAG IPs from untransduced SK-N-MC Tau-GFP KI cell extracts (Fig. 1E & Fig. 3C). PPP2CA is the catalytic subunit of the PP2A holoenzyme complex consisting of the regulatory A and substrate receptor B subunits^51,74–77^. To explore whether the FLAG-aGFP_6M_-PPP2CA and FLAG-aGFP_6M_-PPP2CA^H118Q^ AdPhosphatases transduced in cells could form the full holoenzyme complex and to assess what else they interacted with, we undertook a proteomic analysis on anti-FLAG IPs. As expected, Tau-GFP was one of top interactors of both FLAG-aGFP_6M_-PPP2CA and FLAG-aGFP_6M_-PPP2CA^H118Q^ AdPhosphatases (Table 4). When we analysed the differences, notably, components of the PP2A holoenzyme complex including B subunits PPP2R2A, PPP2R5C, and PPP2R5D, and the A subunit PPP2R1A were enriched only in IPs from FLAG-aGFP_6M_-PPP2CA but not FLAG-aGFP_6M_-PPP2CA^H118Q^ (Fig. 3D), indicating that the catalytically dead PPP2CA^H118Q^ mutant is incapable of forming the full PP2A holoenzyme complex. This reduced interaction of the H118Q mutant with holoenzyme subunits could potentially arise either from the loss of catalytic activity or from mutation-induced conformational changes, although our current data do not distinguish between these possibilities. Additionally, we observed a significantly elevated binding of PPME1 in the FLAG-aGFP_6M_-PPP2CA IPs (Fig. 3D). PPME1 is recognized for its role in negatively regulating PP2A activity ^78^, indicating potential regulation of PP2A activity upon expression of FLAG-aGFP_6M_-PPP2CA AdPhosphatase. We also observed an enrichment of IQGAP1, IQGAP2, MTREX, CAMSAP3 and NUP153 in aGFP_6M_-PPP2CA IPs. It has been reported that PP2A interacts with IQGAP1 to maintain cell-cell adhesion ^79^. Finally, we also conducted phospho-proteomic analysis using data-dependent acquisition mode on anti-GFP IPs from untransduced control SK-N-MC Tau-GFP KI cell extracts or those transduced with FLAG-aGFP_6M_ control, FLAG-aGFP_6M_-PPP2CA or the FLAG-aGFP_6M_-PPP2CA^H118Q^ mutant (Fig. 3E). Through this analysis, we identified 15 phosphorylation sites on TAU that were specifically dephosphorylated by the aGFP_6M_-PPP2CA AdPhosphatase compared to controls (Fig. 3E, Table 5).

### A tractable AdPhosphatase system for targeted dephosphorylation of Tau-GFP

The aGFP_6M_-PPP2CA AdPhosphatase we have used thus far relies on constitutive expression, resulting in a constitutive dephosphorylation of Tau-GFP. To adapt this system to an inducible one, we subcloned aGFP_6M_-PPP1CA and aGFP_6M_-PPP2CA AdPhosphatases into tetracycline-inducible retroviral vectors. Transduction of SK-N-MC Tau-GFP KI cells with retroviruses encoding Tet-inducible expression of aGFP_6M_ control, aGFP_6M_-PPP1CA, aGFP_6M_-PPP1CA^H125Q^ (catalytically inactive mutant), aGFP_6M_-PPP2CA or aGFP_6M_-PPP2CA^H118Q^ led to the expression of these AdPhosphatases only upon doxycycline treatment (Fig. 4A). Excitingly, under these conditions, doxycycline treatment resulted in a decrease in levels of Tau phosphorylation at S396, S181, S404 and S202/T205 but not total Tau in cells transduced with aGFP_6M_-PPP1CA and aGFP_6M_-PPP2CA but not aGFP_6M_ control, aGFP_6M_-PPP1CA^H125Q^, or aGFP_6M_-PPP2CA^H118Q^ (Fig. 4A). Our data not only demonstrate that the AdPhosphatase system can be adapted for an inducible targeted dephosphorylation but also show that the PPP1CA AdPhosphatase is also capable to targeted dephosphorylation of Tau to a similar extent as PPP2CA AdPhosphatase. As expected, anti-FLAG IPs from doxycycline-treated extracts expressing aGFP_6M_, aGFP_6M_-PPP1CA, aGFP_6M_-PPP1CA^H125Q^, aGFP_6M_-PPP2CA or aGFP_6M_-PPP2CA^H118Q^ completely co-precipitate Tau-GFP (Fig. 4B&C), suggesting full target engagement.

**Figure 4:**
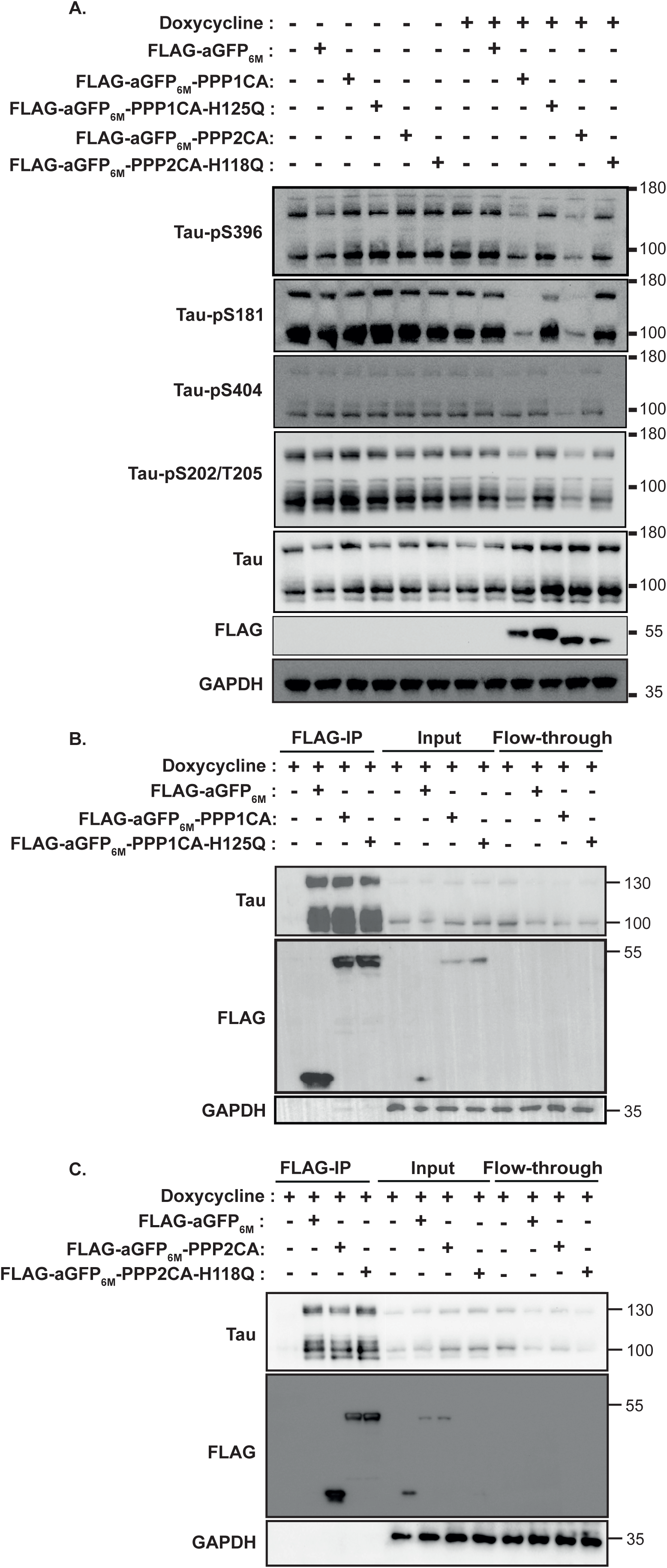
Tractable aGFP_6M_-AdPhosphatases system for targeted dephosphorylation of Tau-GFP. **(A)** *MAPT^GFP/GFP^* KI SK-N-MC cells stably expressing FLAG-aGFP_6M_, FLAG-aGFP_6M_-PPP1CA, FLAG-aGFP_6M_-PPP1CA^H125Q^, aGFP_6M_-PPP2CA or FLAG-aGFP_6M_-PPP2CA^H118Q^ under Tet-inducible promoter were treated with doxycycline (100 ng/ml) for 24 h and lysed. Extracts (20 μg protein) were resolved on 4-12% gradient SDS-PAGE and transferred to nitrocellulose membranes, which were subjected to immunoblotting analysis using the indicated antibodies. **(B)** *MAPT^GFP/GFP^* KI SK-N-MC cells stably expressing FLAG-aGFP_6M_, FLAG-aGFP_6M_-PPP1CA, or FLAG-aGFP_6M_-PPP1CA^H125Q^ under Tet-inducible promoter were treated with doxycycline (100 ng/ml) for 24 h and lysed. Extracts (200 μg protein) were subjected to IP with FLAG-M2 beads. Pre-IP input or post-IP flowthrough extracts (30 μg protein) or FLAG IPs were resolved on 4-12% gradient SDS-PAGE and transferred to nitrocellulose membranes, which were subjected to immunoblotting analysis using the indicated antibodies. **(C)** As in (B) except *MAPT^GFP/GFP^* KI SK-N-MC cells stably expressing FLAG-aGFP_6M_, aGFP_6M_-PPP2CA or FLAG-aGFP_6M_-PPP2CA^H118Q^ under Tet-inducible promoter were used. All blots are representative of at least 3 independent biological replicates.

### AdPhosphatase-mediated dephosphorylation of phospho-Tau-GFP enhances its co-localization and association with microtubules

Tau is a microtubule binding protein, and its phosphorylation is linked to its detachment from microtubules ^7,80–82^. Dephosphorylated Tau displays an increased propensity to associate with microtubules ^83,84^. We therefore tested the impact of AdPhosphatase-mediated targeted dephosphorylation of Tau-GFP on its co-localisation with microtubules in SK-N-MC Tau-GFP KI cells. In untransduced control SK-N-MC Tau-GFP KI cells or those transduced FLAG-aGFP_6M_ or FLAG-aGFP_6M_-PPP2CA^H118Q^ controls, where Tau-GFP is basally phosphorylated (Fig. 1D), low but comparable levels of Tau-GFP signal was detected on microtubules (Fig. 5A&B). In contrast, in SK-N-MC Tau-GFP KI cells transduced with FLAG-aGFP_6M_-PPP2CA, where Tau-GFP is targeted for a robust dephosphorylation on up to 17 phospho-sites (Fig. 3C&E), significantly more Tau-GFP signal was detected on microtubules compared to FLAG-aGFP_6M_-PPP2CA^H118Q^ and other controls (Fig. 5A&B), suggesting that targeted dephosphorylation of Tau leads to its co-localization with microtubules. Next, we performed Tau-GFP immunoprecipitation (IP) followed by mass spectrometry analysis to see if microtubules were co-precipitated. Indeed, we observed an increased abundance of TUBA1B, TUBB and TUBB4B tubulin isoforms co-precipitating with Tau-GFP from cells expressing aGFP6M-PPP2CA compared to controls (untransduced, aGFP_6M_ or aGFP_6M_-PPP2CA^H118Q^) (Fig. 5C). Importantly, the increased co-precipitation of tubulin with dephosphorylated Tau-GFP was also confirmed by immunoblotting of the Tau-GFP IPs with α-tubulin and β-tubulin antibodies (Fig. 5D). Pronounced dephosphorylation of Tau-GFP pSer202/pSer206 by aGFP_6M_-PPP2CA but not by controls (aGFP_6M_ or aGFP_6M_-PPP2CA^H118Q^) is evident in the inputs and Tau-GFP IPs, while flow-through extracts reveal an efficient immune-depletion of Tau-GFP from all samples following anti-GFP IPs, as detected by both anti-Tau and anti-GFP antibodies (Fig. 5D). These observations demonstrate that targeted dephosphorylation of Tau-GFP enhances its co-localization and association with microtubules.

**Figure 5:**
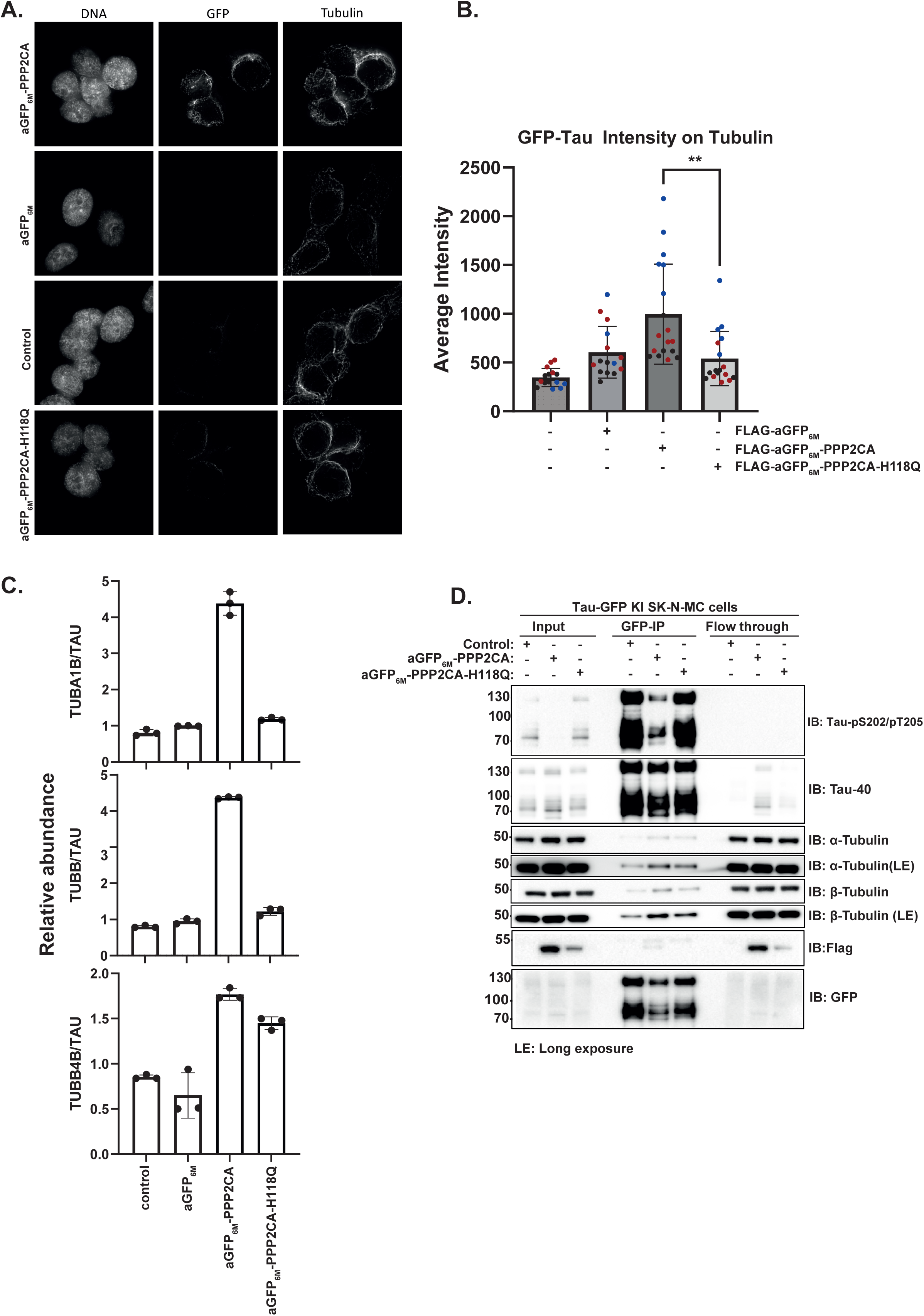
aGFP_6M_-PPP2CA AdPhosphatase-mediated targeted dephosphorylation of Tau-GFP enhances microtubule binding. **(A)** *MAPT^GFP/GFP^* KI SK-N-MC cells (WT) or those stably expressing FLAG-aGFP_6M_, FLAG-aGFP_6M_-PPP2CA, or FLAG-aGFP_6M_-PPP2CA^H118Q^ AdPhosphatases were subjected to pre-extraction of soluble Tau followed by immunofluorescence staining with anti-tubulin antibody. Nucleus was stained with DAPI. Representative images from 3 independent biological replicates are included (n=17). **(B)** The intensity of the GFP fluorescence on images from (A) (n=17 from 3 biological replicates) were measured and average intensity calculated and plotted. Each dot represents average fluorescence intensity on tubulin from a single field of view, and each colour represents a particular biological replicate. Statistical significance was assessed using an unpaired two-tailed t-test. **(C)** SK-N-MC Tau-GFP knock-in (KI) cells transduced with retroviruses encoding the expression of expressing the aGFP_6M_, aGFP_6M_-PPP2CA-FLAG, or catalytically inactive aGFP_6M_-PPP2CA^H118Q^ mutant AdPhosphatases were lysed. Tau-GFP was immunoprecipitated using GFP-Trap beads and subjected to trypsin digestion followed by mass spectrometry analysis. The figure shows the relative abundance of different tubulin isoforms co-precipitated with Tau, represented as the ratio of tubulin isoform intensity to Tau intensity. **(D)** Similar to (C), the indicated cells were lysed and extracts subjected to Tau-GFP immunoprecipitation using GFP-trap beads. Input extracts (10 μg protein), Tau-GFP IPs or post-IP flow through extracts (10 μg protein) were resolved on 4-12% gradient SDS-PAGE and transferred onto nitrocellulose membranes, which were processed by immunoblotting with the indicated antibodies. Almost complete immune-depletion of Tau-GFP from flow-through extracts compared to input demonstrates the efficiency of Tau-GFP immunoprecipitation. The data is representative 3 independent biological replicates.

### Development of small heterobifunctional molecules for proximity-induced dephosphorylation of Tau

The AdPhosphatase system described above relies on retroviral transduction and expression of the aGFP_6M_-AdPhosphatases, which permanently associate with Tau-GFP in cells. An ideal targeted dephosphorylation system would involve inducing transient proximity between an endogenous phosphatase and the target phospho-protein with a small cell-permeable molecule. For proof-of-concept for such a system, we developed a small heterobifunctional molecule with established ligands recognising bromoTAG ^85^ on one end and FKBP12^F36V^(dTAG) ^86^ on the other, with each ligand connected through 3-polyethylene glycol (PEG) linker (Fig. 6A). The molecule was named BDPIC (bromoTAG-dTAG proximity-inducing chimera). By using a cell-based NanoBRET ternary complex assay ^87^, when we transiently overexpressed dTAG-NanoLuc and bromoTAG-HALO in U2OS osteosarcoma cells, at 200 nM BDPIC treatment induced a robust bioluminescence resonance energy transfer compared to DMSO treatment (Fig. 6B), suggesting that BDPIC induces a robust interaction between dTAG and bromoTAG in cells. The NanoBRET assay was optimised in U2OS cells with different orientations of dTAG-NLuc and bromoTAG-HALO or dTAG-HALO and bromoTAG-NLuc constructs transiently overexpressed to determine the optimal combination for a NanoBRET ternary complex formation (Fig. S6A-B). For the optimal combination (dTAG-NanoLuc and bromoTAG-HALO), BDPIC induced a dose-dependent increase in bioluminescence resonance energy transfer from 10-200 nM BDPIC compared to DMSO or no ligand control. However, the signal was slightly lower at 1000 nM compared to 200 nM (Fig. S6C), potentially implying a well-known hook effect for heterobifunctional molecules at high concentrations^88^. After establishing that BDPIC can induce proximity between dTAG and bromoTAG in cells, we sought to investigate whether applying different doses of BDPIC in cells expressing bromoTAG-PPP2CA and Tau-dTAG would reveal an optimal dose which can induce proximity between them and allow bromoTAG-PPP2CA to dephosphorylate Tau-dTAG. To achieve this, we generated Tau-dTAG knock-in SK-N-MC cells using CRISPR/Cas9 genome editing, by incorporating *FKBP12^F36V^* on *MAPT* alleles, and validated these by immunoblotting and genomic DNA sequencing (Fig. S7A-C). In Tau-dTAG KI SK-N-MC cells stably expressing HA-bromoTAG-PPP2CA, compared to DMSO-treated controls, BDPIC treatment at both 100 nM and 250 nM for 24 h resulted in a robust Tau-dTAG dephosphorylation at S396, S202/T205, S181 and S404 (Fig. 7C). In contrast, in cells expressing empty vector, HA-bromoTAG, or catalytically inactive HA-bromoTAG-PPP2CA^H118Q^ mutant, BDPIC treatment did not alter the levels of Tau-dTAG phosphorylation at S396, S202/T205, S181 and S404 compared to DMSO-treated controls (Fig. 7C). Since the targeted dephosphorylation caused by BDPIC is through induced proximity, we reasoned that swapping the bromoTAG and dTAG tags on Tau and PPP2CA should still allow BDPIC to mediate targeted dephosphorylation of Tau. Hence, we generated homozygous Tau-bromoTAG KI SK-N-MC cells using CRISPR/Cas9 genome editing and validated these by immunoblotting (Fig. S7D-F). In these cells, when FLAG-dTAG-PPP2CA was stably expressed, BDPIC treatment at 100 nM, 250 nM or 500 nM for 24 h resulted in a robust Tau-bromoTAG dephosphorylation at S396, and S202/T205 compared to DMSO-treated controls (Fig. 6D). However, BDPIC treatment did not cause any Tau-bromoTAG dephosphorylation in cells stably expressing empty vector or FLAG-dTAG-PPP2CA^H118Q^ compared to DMSO treatment (Fig. 6D). These data demonstrate that BDPIC-induced proximity between catalytically active PPP2CA, but not the inactive mutant, and Tau induces targeted dephosphorylation of Tau. The observation that BDPIC facilitates the targeted dephosphorylation of TAU-dTAG by HA-bromoTAG-PPP2CA and TAU-bromoTAG by FLAG-dTAG-PPP2CA suggests the formation of a ternary complex. Indeed, anti-FLAG IPs from Tau-bromoTAG KI SK-N-MC cells stably expressing FLAG-dTAG-PPP2CA or FLAG-dTAG-PPP2CA^H118Q^ both co-precipitated Tau-bromoTAG upon BDPIC but not DMSO treatment (Fig. 6E).

**Figure 6:**
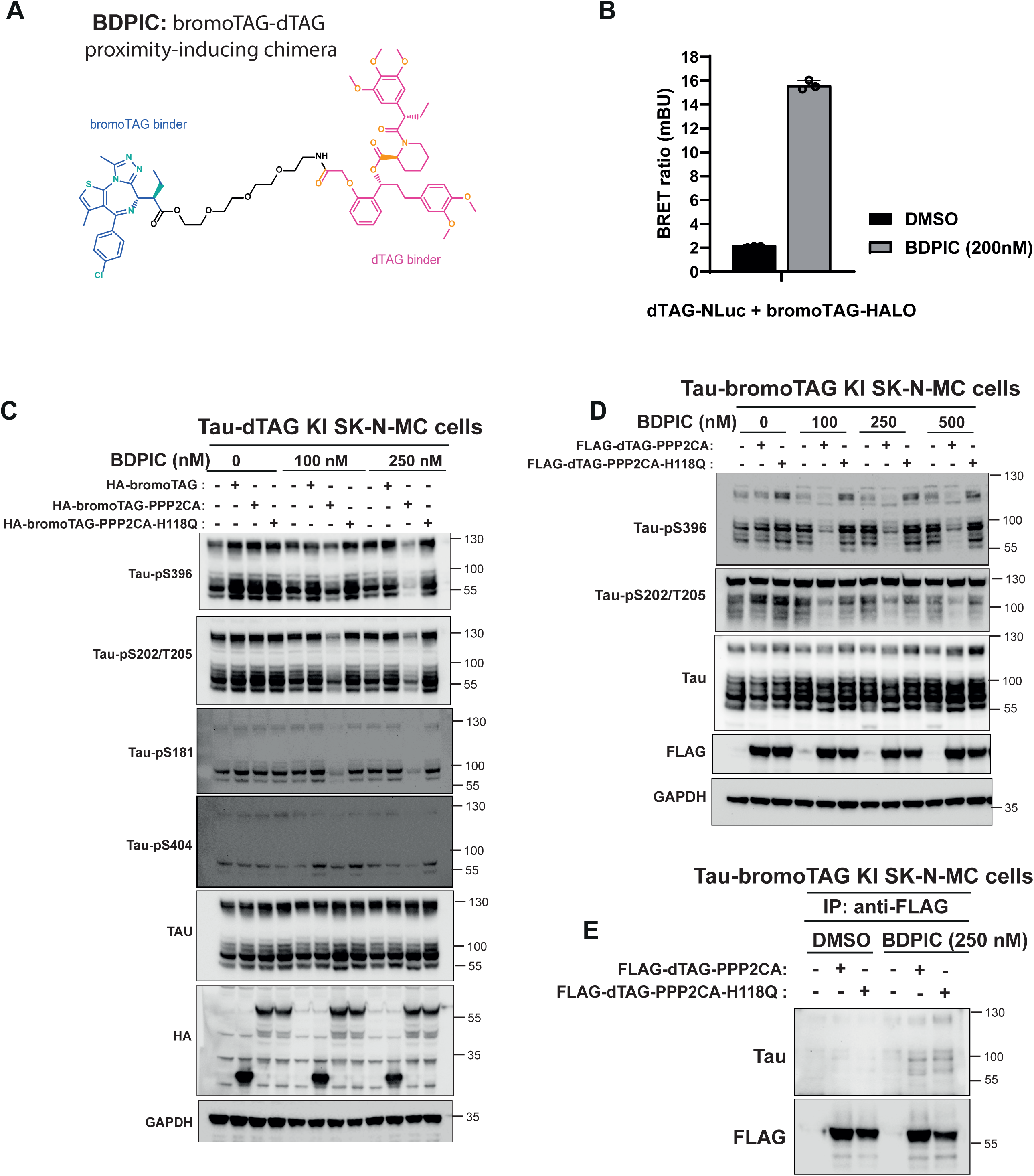
Targeted dephosphorylation of endogenously knocked in Tau-dTAG or Tau-bromoTAG via BDPIC-induced recruitment of bromoTAG-PPP2CA and dTAG-PPP2CA, respectively. **(A)** Structure of the heterobifunctional BDPIC (bromoTAG-dTAG proximity-inducing chimera) molecule with Brd4BD2^L387A^ (bromoTAG)- and FKBP12^F36V^ (dTAG)-binding warheads connected with a 3XPEG linker. **(B)** A NanoBRET ternary complex formation assay was performed in U2OS cells that were transiently transfected with the vectors encoding dTAG-NLuc and bromoTAG-HALO. Cells were plated onto white 96-well plates, treated with DMSO or BDPIC at 200 nM for 24 h. BRET signal ratios were determined and plotted in the graph (n=3). **(C)** Tau-dTAG KI SK-N-MC cells or those stably expressing HA-bromoTAG, HA-bromoTAG-PPP2CA or HA-bromoTAG-PPP2CA^H118Q^ were treated with 0, 100 and 250 nm BDPIC for 24 h prior to lysis. Extracts (20 μg protein) were resolved on 4-12% gradient SDS-PAGE and transferred to nitrocellulose membranes, which were subjected to immunoblotting analysis using the indicated antibodies. **(D)** As in (C) except *MAPT^bromoTAG/bromoTAG^* KI SK-N-MC cells or those stably expressing FLAG-dTAG-PPP2CA or FLAG-dTAG-PPP2CA^H118Q^ were treated with 0, 100, 250 and 500 nm BDPIC for 24 h prior to lysis. **(E)** *MAPT^bromoTAG/bromoTAG^*KI SK-N-MC cells or those stably expressing FLAG-dTAG-PPP2CA or FLAG-dTAG-PPP2CA^H118Q^ were treated with DMSO or 250 nm BDPIC for 24 h prior to lysis. Extracts were subjected to anti-FLAG IPs. The IPs were resolved on 4-12% SDS-PAGE, transferred to nitrocellulose membranes, which were subjected to immunoblotting analysis using the indicated antibodies. All blots are representative of at least 3 independent biological replicates.

**Figure 7:**
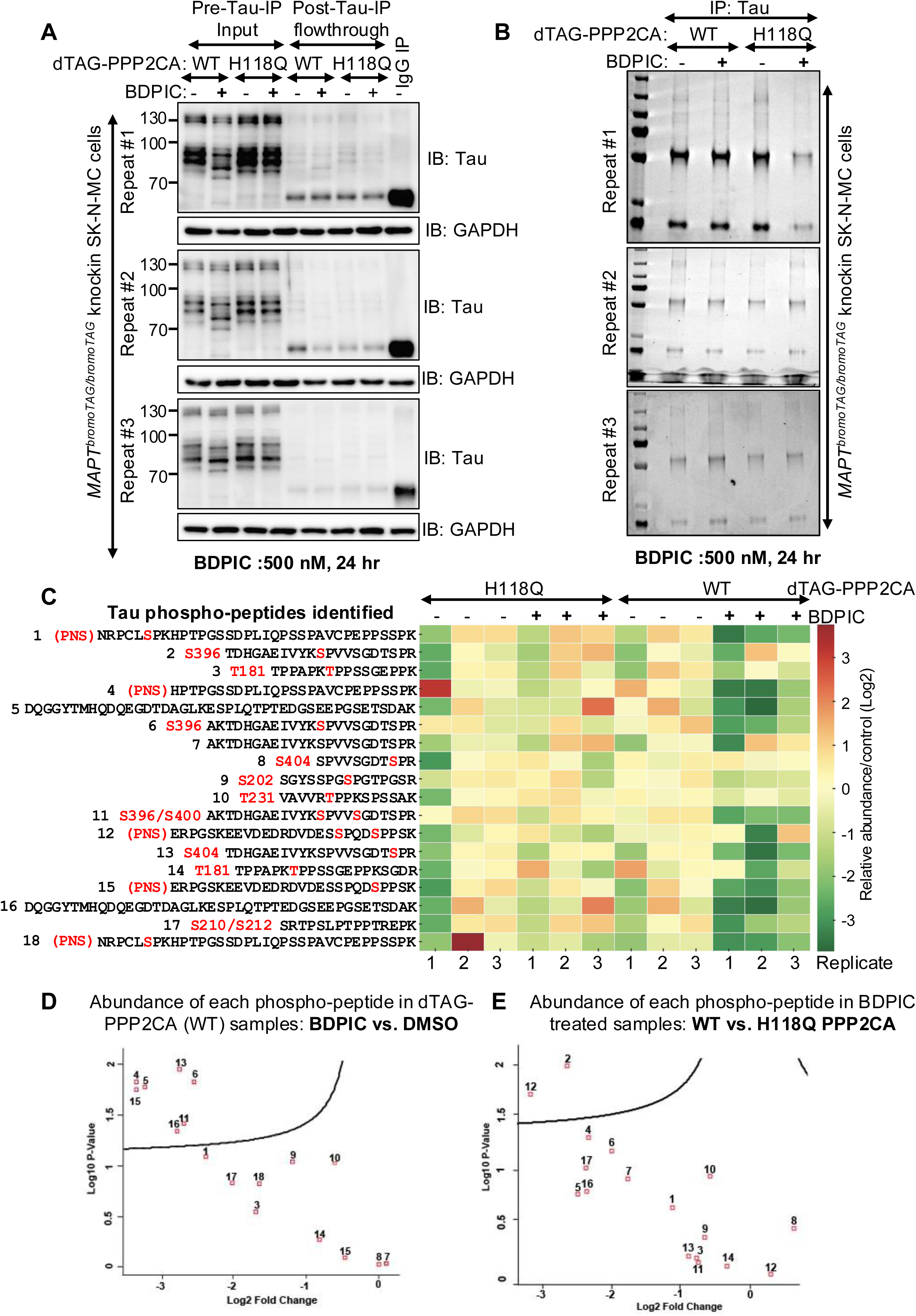
Proteomic analysis of the BDPIC-mediated targeted dephosphorylation of Tau-bromoTAG via dTAG-PPP2CA. **(A)** *MAPT^bromoTAG/bromoTAG^* KI SK-N-MC cells stably expressing FLAG-dTAG-PPP2CA or FLAG-dTAG-PPP2CA^H118Q^ were treated with BDPIC for 24 h prior to lysis. Tau-bromoTAG extracts (1.5 mg) were subjected to immunoprecipitation using anti-Tau antibody. Pre-IP input and post-IP flowthrough extracts (20 μg protein) were resolved on 4-12% gradient SDS-PAGE and transferred to PVDF membranes, which were subjected to immunoblotting analysis using the indicated antibodies. All 3 biological replicates are included. Interestingly, a change in the mobility shift downwards of all Tau variants in BDPIC-treated cells expressing dTAG-PPP2CA compared to DMSO-treated controls is visible, indicating targeted dephosphorylation. **(B)** Tau IPs from (A) were resolved on 4-12% gradient SDS-PAGE and gels stained with Coomassie InstantBlue. Boxed gel pieces were excised and processed for in-gel tryptic digestion and subsequent phospho-proteomic analysis. **(C)** Tryptic peptides from (B) were analysed using data dependent analysis (DDA). The heatmap represents the relative abundance of the identified phospho-Tau peptides across each condition and biological replicate. Where it was possible to unequivocally identify the phospho-site within the identified peptide, the residue number corresponds to 2N4R variant of Tau. PNS – peptide from PNS Tau variant. **(D)** A volcano plot comparing the fold-change in abundance of phospho-peptides identified in (C) in *MAPT^bromoTAG/bromoTAG^* KI SK-N-MC cells stably expressing FLAG-dTAG-PPP2CA treated with BDPIC against DMSO control. 7 phospho-peptides are significantly downregulated by BDPIC treatment compared to DMSO controls. **(E)** A volcano plot comparing the fold-change in abundance of phospho-peptides identified in (C) in BDPIC-treated *MAPT^bromoTAG/bromoTAG^* KI SK-N-MC cells stably expressing FLAG-dTAG-PPP2CA vs. FLAG-dTAG-PPP2CA-H118Q. 2 phospho-peptides are significantly downregulated by BDPIC treatment in cells expressing dTAG-PPP2CA compared to those expressing dTAG-PPP2CA-H118Q.

### Mapping endogenous Tau-bromoTAG phospho-sites targeted for dephosphorylation via BDPIC-mediated recruitment of dTAG-PPP2CA

To gain deeper insight into the extent of targeted dephosphorylation of Tau-bromoTAG by BDPIC via dTAG-PPP2CA, we undertook a mass spectrometry analysis to identify changes in Tau phospho-sites upon targeted dephosphorylation by BDPIC. For this, Tau-bromoTAG KI SK-N-MC cells stably expressing dTAG-PPP2CA or dTAG-PP2CA^H118Q^ mutant were treated with DMSO or BDPIC in 3 biological replicates (Fig. 7A). As evident in total Tau immunoblots, in Tau-bromoTAG KI SK-N-MC cells stably expressing dTAG-PPP2CA, but not in those expressing dTAG-PP2CA^H118Q^, BDPIC (500 nM for 24h) treatment causes a downward mobility shift of all Tau variants compared to DMSO treated controls (Fig. 7A), potentially suggesting extensive targeted dephosphorylation of Tau. From these extracts, endogenous Tau-bromoTAG was immunoprecipitated with an anti-Tau antibody, and the complete depletion of Tau signal from post-IP flowthrough extracts compared to pre-IP input extracts demonstrated an efficient capture of Tau-bromoTAG by anti-Tau IPs (Fig. 7A). Tau-bromoTAG IPs were resolved via SDS-PAGE and all areas where Tau variants migrate were excised and subjected to in-gel tryptic digestion (Fig. 7B). The resulting peptides were isolated and analysed by mass spectrometry. The data analysis was conducted using the SEQUEST search engine by filtering for phospho-peptides corresponding to Tau and normalized to total Tau protein intensity. Statistical analysis was performed using Perseus. From 3 biological replicates, a total of 18 Tau phospho-peptides were identified. In cells stably expressing dTAG-PPP2CA and treated with BDPIC, a reduction in phosphorylation was observed at most Tau phospho-sites identified compared to DMSO-treated controls, with 7 out of 18 phospho-sites showing significant dephosphorylation (Fig. 7C&D, Table 6). Similarly, in cells treated with BDPIC, those stably expressing dTAG-PPP2CA resulted in reduction in phosphorylation at most Tau phospho-sites compared to those stably expressing the dTAG-PP2CA^H118Q^ mutant, with 2 sites exhibiting significant dephosphorylation (Fig. 7C&E, Table 7). The heatmap and volcano plots of all 18 phospho-peptides show their relative abundance across different conditions (Fig. 7C-E). The Tau phospho-sites that are dephosphorylated by BDPIC by recruiting dTAG-PPP2CA are shown along the domain architectures of the 2N4R Tau variants (Fig. S8).

### Proof-of-concept for BDPIC-mediated dephosphorylation of phospho-TAU-bromoTAG by redirecting dTAG-PPP2CA at the endogenous level

Thus far, we have assessed the targeted dephosphorylation of Tau at the endogenous level by redirecting phosphatases that were exogenously introduced in cells. To assess whether PPP2CA at the endogenous level could be redirected to dephosphorylate Tau, we knocked-in *dTAG* on *PPP2CA* gene in Tau-bromoTAG KI SK-N-MC cells using CRISPR/Cas9 genome editing technology and obtained several heterozygous *^dTAG/+^PPP2CA* clones. Knock-ins were confirmed by immunoblotting and DNA sequencing (Fig. S9A-C). Treatment of several clones of the TAU-bromoTAG and dTAG-PPP2CA double KI SK-N-MC cells with increasing concentrations of BDPIC for 24 h elicited a similar dose-dependent reduction in phosphorylation of TAU-bromoTAG at S202/T205 compared to DMSO-treated controls, with maximal dephosphorylation observed with 250 nM BDPIC (Fig. 8A, Fig. S9D), with a slight rescue of phosphorylation observed at 1000 nM BDPIC, reminiscent of a hook-effect reported for bivalent PROTAC molecules at high concentrations ^89^. Treatment of the parental Tau-bromoTAG KI SK-N-MC cells harbouring unmodified PPP2CA with BDPIC showed similar levels of phosphorylation of TAU-bromoTAG at S202/T205 compared to DMSO controls (Fig. 6D, S7D). The data indicates that BDPIC-induced proximity between endogenous dTAG-PPP2CA and Tau-bromoTAG can effectuate targeted dephosphorylation of Tau. When Tau-bromoTAG and dTAG-PPP2CA double KI SK-N-MC cells were treated with 250 nM BDPIC over a period of 24 h, robust Tau-bromoTAG dephosphorylation at S202/T205 was observed within 30 minutes, while maximal dephosphorylation was observed as early as 2 h, which was sustained over 24 h following BDPIC treatment (Fig. 8B). We also conducted competition assays using excess amounts of dTAG^V^-1-NEG (inactive enantiomer of the dTAG-V1 PROTAC that shares the same dTAG binder as BDPIC but is unable to recruit VHL for dTAG degradation) ^90^ and cis-AGB1 (an inactive enantiomer of the AGB1 PROTAC that shares the same bromoTAG binder as BDPIC but cannot recruit VHL to degrade the bromoTAG protein)^85^. Pre-treatment of cells with excess dTAG^V^-1-NEG or cis-AGB1 prevented the dephosphorylation of Tau-bromoTAG induced by BDPIC in TAU-bromoTAG and dTAG-PPP2CA double knock-in SK-N-MC cells (Fig. 8C). These data confirm that BDPIC-mediated targeted dephosphorylation of Tau-bromoTAG relies on the binding of BDPIC to both Tau-bromoTAG and dTAG-PPP2CA.

**Figure 8:**
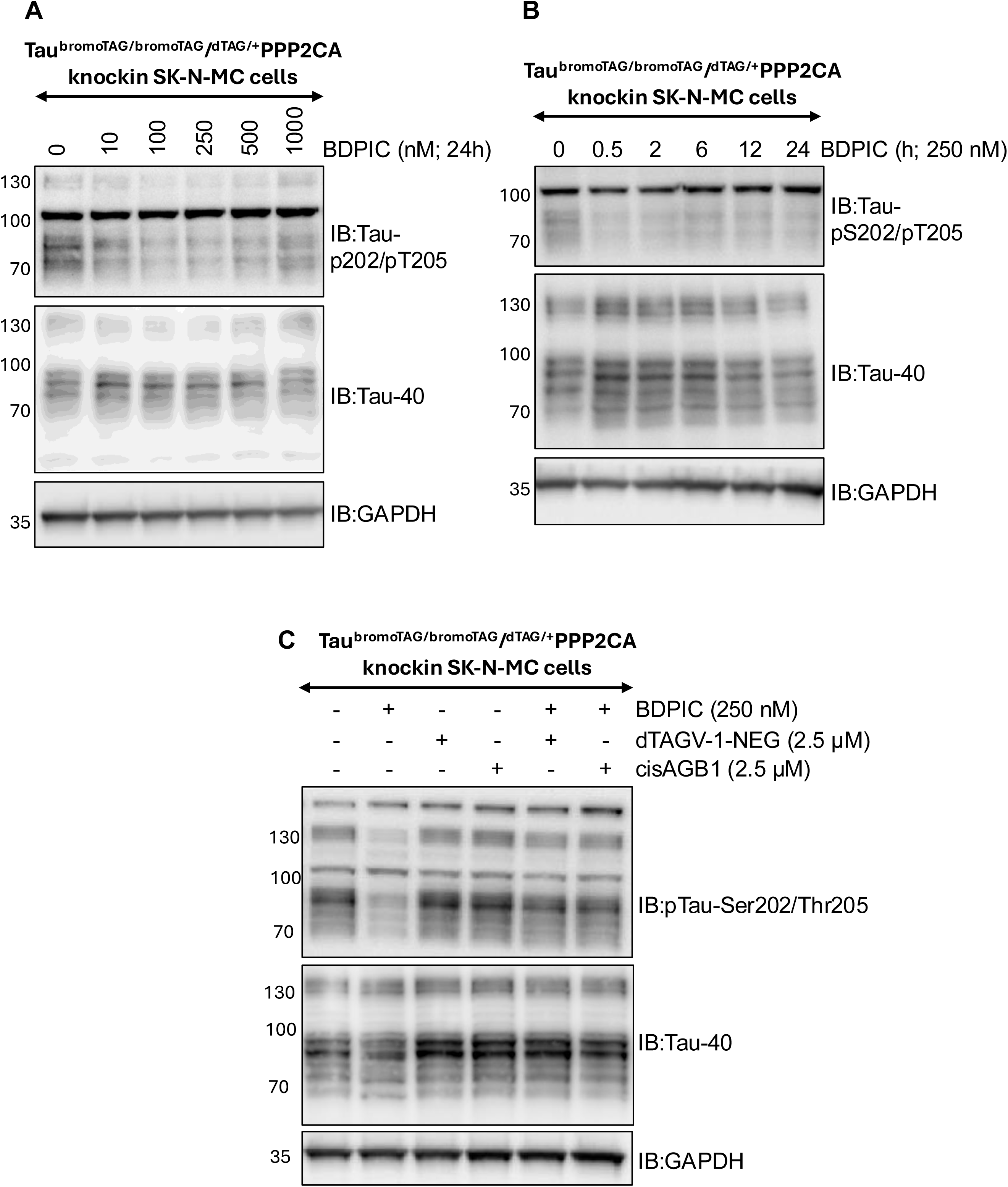
BDPIC-induced targeted dephosphorylation of TAU-bromoTAG via recruitment of dTAG-PPP2CA at the endogenous level. **(A)** *MAPT^bromoTAG/bromoTAG^/^+/dTAG^PPP2CA* double KI SK-N-MC cells (Cl3) were generated by CRISPR/Cas9 technology. Cells were treated with increasing concentration of BDPIC (0-1000 nM) for 24 h prior to lysis. Extracts (25 μg protein) were resolved on 4-12% gradient SDS-PAGE and transferred to PVDF membranes, which were subjected to immunoblotting analysis using the indicated antibodies. **(B)** Cells from (A) were treated with BDPIC (250 nM) for a time course experiment as indicated from 0-24 h prior to lysis. Extracts (25 μg protein) were resolved on 4-12% gradient SDS-PAGE and transferred to PVDF membranes, which were subjected to immunoblotting analysis using the indicated antibodies. **(C)** As in (B) except *MAPT^bromoTAG/bromoTAG^/^+/dTAG^PPP2CA* double KI SK-N-MC cells were treated with DMSO or BDPIC (100 nM) in the presence or absence of excess cis-AGB1 and dTAG^V^-1 NEG (2.5 μM) for 24 h prior to lysis. All blots are representative of at least 3 independent biological replicates.

## Discussion

Despite advances in our understanding of the causes and features of tauopathies, such as AD, there are currently no effective disease-modifying therapies ^91^. Thus, there is a pressing need for innovative therapeutic approaches against tauopathies. In this study, we explored targeted dephosphorylation as a strategy to modulate Tau function at the endogenous level. By redirecting aGFP_6M_-PPP1CA and aGFP_6M_-PPP2CA AdPhosphatases to Tau-GFP, which was homozygously knocked in at the *MAPT* locus in SK-N-MC neuroblastoma cells, we achieved targeted dephosphorylation of Tau at multiple sites, including S46 ^92^, S516 ^35^, S519 ^93^, T529 ^49^, T548 ^94^, S673 ^95^, S713 ^35^, S717 ^96^, and T720 ^97^ that have been reported to play important roles in AD pathogenesis. Excitingly, the targeted dephosphorylation of Tau enhanced its microtubule binding. These findings provide proof-of-concept that our AdPhosphatase system effectively targets the dephosphorylation of phospho-sites on Tau that not only regulate microtubule binding but are also important for driving AD pathogenesis. Moreover, we demonstrated proof-of-concept for small bivalent molecule-induced dephosphorylation of Tau at the endogenous level by using the BDPIC (bromoTAG-dTAG proximity-inducing chimera) system. In CRISPR-engineered cells harbouring dTAG-PPP2CA and Tau-bromoTAG double knock-ins, BDPIC-induced dephosphorylation of Tau on multiple phospho-sites via the recruitment of PPP2CA at the endogenous level. Indeed, through a phospho-proteomic analysis in endogenous Tau-bromoTAG IPs, we were able to identify 5 phospho-sites (pT181, pS202, pT231, pS396 and pS404) that were significantly dephosphorylated by BDPIC-induced recruitment of dTAG-PPP2CA catalytic activity. 4 out of these 5 phospho-sites have been reported to play critical roles in driving Tau aggregation and AD pathology ^28–30,35^. Furthermore, of the 7 more phospho-sites that were also identified, most also showed reduced abundance following BDPIC-induced recruitment of dTAG-PPP2CA catalytic activity to Tau-bromoTAG, although the reduction was not significant. Although the AdPhosphatase system demonstrated that targeted recruitment of PPP2CA can induce selective dephosphorylation of endogenous Tau, it relies on genetic delivery, and the phosphatase is constantly bound to Tau. To circumvent these limitations, we developed the BDPIC heterobifunctional small-molecule approach, where we achieve chemically inducible, fully reversible and pharmacologically tractable control of Tau dephosphorylation. Together, these complementary genetic and chemical proximity-based strategies establish the mode of action and demonstrate the feasibility of recruiting endogenous phosphatases to POIs for targeted dephosphorylation. Given the role of Tau hyperphosphorylation in AD, we propose targeted dephosphorylation via small molecules as a promising therapeutic strategy against AD and other tauopathies. However, Tau phosphorylation is dynamic, and is influenced by neuronal activity and localization ^11^. While most studies imply that hyperphosphorylation of Tau promotes aggregation and Tau pathology, it is likely that some phosphorylation may be neuroprotective. Therefore, indiscriminate targeted Tau dephosphorylation could potentially risk leading to unwanted side-effects, especially if neuroprotective phospho-sites are targeted. Future work should map the tau phospho-landscape by disease staging and cell types, identify critical pathogenic phospho-epitopes, and develop tools to deliver phosphatases to precisely dephosphorylate these sites. Furthermore, combining Tau dephosphorylation with Tau clearance via degraders, aggregation inhibitors, or microtubule stabilizers may enhance efficacy.

Few other studies have recently reported targeted dephosphorylation of Tau. By using a cell permeable bivalent peptides consisting of Tau and PP1 & PP2A-Bα binders, it was shown that the peptide caused a reduction in levels of both phospho- and total Tau ^83,98,99^. Given this synergistic reduction, these studies failed to demonstrate definitive evidence for targeted dephosphorylation versus degradation of Tau ^83,98^. In another study developing proof-of-concept for targeted dephosphorylation using small bivalent molecule PhosTAC7, a molecule which was designed to simultaneously recruit dTAG and HaloTag in cells, a reduction in phosphorylation levels of overexpressed Halo-tagged Tau was shown in cells co-expressing the regulatory subunit of PP2A, dTAG-PP2A-A, although the effect of the compound in the absence of dTAG-PP2A-A was not tested ^100^. This would be important control, as in our assays PhosTAC7 and another dTAG-HaloTag proximity-inducing chimera (HDPIC) failed to recruit dTAG-phosphatases to Halo-tagged target phospho-proteins of interest (POIs) for dephosphorylation, and instead the molecules appeared to destabilize Halo-tagged POIs through binding to the chloroalkane moiety independent of dTAG-binding ^54^. A series of bifunctional molecules consisting of PET-probes targeting Tau and a dTAG binder have also been reported to cause a slight reduction in phosphorylation of overexpressed Tau in cells also overexpressing dTAG-PP2A-A ^100^, even though the PET tracers are reported to only bind aggregated Tau neurofibrillary tangles. While these studies focussed on the impact of targeted dephosphorylation on overexpressed Tau, none explored targeted dephosphorylation of Tau and its function at the endogenous levels. Furthermore, none of these studies fully established the requirement of the recruited phosphatase activity for the reduction in Tau phosphorylation that was observed ^83,98–100^.

A limitation of this study is that the endogenous abundance of PPP2CA was not systematically assessed across the cell lines in which we assessed the abundance of Tau. Although PPP2CA is generally highly and ubiquitously expressed, cell-specific differences in its abundance, localization or holoenzyme composition may influence the efficiency of heterobifunctional molecule-induced Tau dephosphorylation. Future studies will focus on targeted dephosphorylation of Tau in aggregate models to interrogate whether dephosphorylation of Tau impedes its propensity to aggregate and cause cytotoxicity. We are also working to develop protein phosphatase binders with the hope of being able to convert them into bivalent phosphatase-recruiting chimeras (PhosTACs). Ultimately, to convert proof-of-concept ideas into potential drugs that effectuate targeted dephosphorylation *in vivo*, we need robust phosphatase ligands that allow us to hijack endogenous phosphatases.

## Acknowledgements

GPS is funded by the UKRI Medical Research Council (MC_UU_00018/6 and MC_UU_00038/6) and is also supported by Boehringer-Ingelheim through the Division of Signal Transduction Therapy (University of Dundee). GS & TC are supported by UKRI MRC funds awarded to GPS. FB is supported by UKRI MRC PPU PhD studentship. J-FZ is supported by the International Alliances between University of Copenhagen and University of Dundee using the funding from the Novo Nordisk Foundation (Grant number NNF18CC0034900 and NNF23SA0084103). RS is funded by grant award from Amgen Inc. made jointly to Alessio Ciulli, Adrian Saurin & GPS. We thank the members of GPS lab for experimental advice and discussions during the study. We thank the MRC tissue culture facility staff (E. Allen, A. Muir, S. Dalglish, E. Webster and J. Stark), the staff at the DNA Sequencing services (School of Life Sciences, University of Dundee), the cloning, antibody, inhibitor and protein production teams within the MRC PPU Reagents & Services (University of Dundee) coordinated by J. Hastie and the staff at FACS service for their technical and reagents support. We thank members of the Dundee Imaging Facility (University of Dundee, coordinated by P. Appleton) for microscopy support. We thank the MRC PPU mass spectrometry service team for their help with the project.

## Author Contributions

GS performed most of the experiments, collected and analysed data and contributed to the writing of the manuscript. FB performed IP/MS experiments for BDPIC-induced dephosphorylation of Tau and generated dTAG-PPP2CA/Tau-bromoTAG double KI SK-N-MC cells and performed subsequent experiments. TC undertook IF experiments and collected and analysed data. RS optimised and performed ternary complex assays for BDPIC. J-FZ and ARB performed some experiments and collected and analysed data. TJM, NTW and J-FZ generated constructs used in this study. TJM designed the strategies, and generated constructs, for CRISPR/Cas9 genome editing experiments. GPS conceived and supervised the project, analysed data and contributed to the writing of the manuscript.

## Lead Contact

Further information and requests for resources and reagents should be directed to and will be fulfilled by the Lead Contact, Gopal Sapkota.

## Data and Code Availability

The datasets generated during this study are publicly available at Mendeley Data (https://data.mendeley.com/preview/2r4ys489zp?a=750d63e4-be11-4a28-b95d-680fd666ab83). All raw mass spectrometry data acquired from this study has been deposited in the PRIDE Archive, with the accession number PXD054610 (Phospho-proteome), PXD054554 (total proteome). This paper does not report original code.

## Materials and Methods

### Materials

The following primary antibodies were employed in this study: From Cell Signalling Technology (CST), we used anti-pTau-S404 (Cat# 20194), anti-pTau-S396 (Cat# 9632), anti-pTau-S396 (Cat# AB9658), anti-pTau-S202/T205 (Cat# 30505S), anti-pTau-T181 (Cat# 12885S). Anti-GAPDH-HRP antibody (Cat# HRP-60004), anti-Alpha-Tubulin-HRP conjugated antibody (Cat# HRP-66031) and anti-FLAG M2-HRP antibody (Cat# 86861) were from Proteintech. Anti-β-Tubulin I antibody (Cat# T7816) were from Sigma-Aldrich. Sheep anti-Tau40 (Cat# S157B), sheep anti-dTAG (Cat# DA179), sheep anti-bromoTAG (Cat# SA599), sheep anti-GFP (Cat# S268B) antibodies were generated by the MRC PPU Reagents and Services at University of Dundee. For secondary antibodies, we utilized goat anti-Rabbit IgG (H+L)-HRP (#7074), and horse anti-mouse IgG (H+L)-HRP (#7076), both procured from CST. Furthermore, goat anti-Rabbit IgG (H+L)-HRP (#31460), and rabbit anti-sheep IgG (H+L)-HRP (#31480) were purchased from Thermo Fisher Scientific. Eagle’s minimum essential medium (EMEM) was procured from ATCC, and Fetal Bovine Serum (FBS) was obtained from Thermo Fisher Scientific. All plasmids employed in this research were generated by MRC PPU Reagents and Services at University of Dundee, with detailed information accessible at https://mrcppureagents.dundee.ac.uk. BDPIC was synthesized by Natalia Shpiro (Dundee).

### Cell culture

Human embryonic kidney cells 293FT cells were cultured in DMEM with 10% FBS, 2 mM L-glutamine and 1% streptomycin/penicillin. SK-N-MC human neuroblastoma cells were cultured in EMEM with 10% FBS, 2 mM L-glutamine and 1% streptomycin/penicillin. All cell lines were maintained at 37°C and under 5% CO_2_ in a humidified tissue culture incubator. Cells were exposed to different stimuli and compounds as described in the appropriate figure legends prior to lysis. Extracts from the following cells were obtained from members of the Sapkota lab: SH-SY5Y neuroblastoma, HPAF-II human pancreatic adenocarcinoma, HeLa ovarian cancer, MCF-7 breast cancer, T-47D breast cancer, ARPE-19 retinal pigment epithelial, A549 adenocarcinoma, SK-MEL-13 melanoma, DLD-1 colorectal cancer, NCI-H23 adenocarcinoma, HepG2 hepatoma, NCI-H460 lung cancer, U2OS osteosarcoma, HaCaT keratinocytes, HCT-116 colon cancer and Jurkat T-lymphocytes.

### Plasmids

For production of retroviral vectors, the following were cloned into pBabeD-puromycin plasmids: FLAG-aGFP_6M_ (DU60521), FLAG-aGFP_6M_-PPP2CA (DU62902), FLAG-aGFP_6M_-PPP2CA^H118Q^ (DU62960), 3HA-bromoTAG (DU71673), 3HA-bromoTAG-PPP2CA (DU75905), and 3HA-bromoTAG-PPP2CAH118Q (DU75906). All constructs were sequence-verified by the DNA Sequencing Service, University of Dundee (http://www.dnaseq.co.uk). For CRISPR/Cas9 gene editing, the following guide RNAs (gRNA) and donor cDNAs were used: To knockin GFP, dTAG or bromoTAG at the C-terminus of human Tau: sense gRNA (DU74043), antisense gRNA (DU74044), GFP-tag donor (DU74042), dTAG-IRES2-GFP donor (DU74681), bromoTAG-IRES2-mCherry donor (DU74680). To knockin dTAG at the N-terminus of human *PPP2CA* gene, we used the GFP-IRES-dTAG cassette donor and gRNAs as reported previously ^51,53^. All constructs are available to request from the MRC-PPU Reagents and Services webpage (http://mrcppureagents.dundee.ac.uk) and the unique identifier (DU) numbers also provide direct links to the cloning strategies and sequence details.

### Retroviral generation of stable cell lines

Stable cell lines were established via retroviral transduction. The cDNA of interest was introduced into a pBabeD-puromycin vector (6 μg) and co-transfected with pCMV5-GAG/POL (3.2 μg) and pCMV5-VSV-G plasmids (2.8 μg) (Clontech) into a 10-cm dish containing ∼60-70% confluent HEK-293FT cells using PEI reagent as previously described ^52^. 48 hours post-transfection, the medium containing the retroviral particles was collected, filtered through a 0.45 mm filter, and added to target cells along with 10 μg/ml polybrene (Sigma-Aldrich). 48 h later, cells were subjected to selection with 2 μg/ml puromycin (Sigma-Aldrich) to select transduced cells. A pool of transduced cells was then employed for subsequent experiments.

### Cell Lysis

Cells were collected by rinsing twice with phosphate-buffered saline (PBS) and then gently scraped into ice-cold lysis buffer. The lysis buffer composition consisted of 50 mM HEPES pH 7.4, 150 mM NaCl, 1 mM EDTA, 10% glycerol, 0.5% NP-40, 1 mM DTT, 1 mM PMSF, 1.15 mM sodium molybdate, 4 mM sodium tartrate, 10 mM β-glycerophosphate, 1 mM sodium fluoride, 1 mM sodium orthovanadate, and 1x complete protease inhibitor cocktail (Roche). Following a 10-minute incubation on ice, the lysates were clarified by centrifugation at 20,000 x g for 10 minutes at 4°C. The protein concentration was determined using the BCA protein assay.

### Generation of cell lines using CRISPR/Cas9

For the generation of GFP, dTAG or bromoTAG knockin (KI) cells by CRISPR/Cas9 genome editing, previously established protocols were employed ^51,52,101,102^. Briefly, to generate Tau-GFP, Tau-dTAG or Tau-bromoTAG KI cells, SK-N-MC cells were transfected with vectors encoding a pair of guide RNAs (pBabeD-puro-sgRNA1 (1 μg) and pX335-CAS9-D10A-sgRNA2 (1 μg)) targeting close to the stop codon of the *MAPT* gene, along with the respective donor plasmid carrying the in-frame GFP, dTAG-IRES-GFP or bromoTAG-IRES-mCherry KI donors that were flanked by *MAPT* homology arms around the insertion site (3 μg). For the generation of human dTAG-PPP2CA KI cells, *Tau^bromoTAG/bromoTAG^* KI SK-N-MC cells were transfected with vectors encoding a single gRNA (pX459-Puro-CAS9-gRNA (1 μg)) targeting close to the start codon of the *PPP2CA* gene and donor plasmid carrying GFP-IRES2-dTAG KI cassette flanked by *PPP2CA* homology arms (3 μg). One day post-transfection, cells were selected with 2 μg/ml puromycin for a further 48 h. The transfection process was repeated one more time when cells were 60-70% confluent following selection in puromycin. 24 h post-transfection, cells were maintained in fresh medium until single cell isolation by FACS. For the acquisition of single-cell clones, single cells were isolated by fluorescence-activated cell sorting (FACS) using an Influx cell sorter (Becton Dickinson) using either GFP or mCherry channels as appropriate. Negative fluorescence gates were set using non-fluorescent wild type SK-N-MC control cells. Fluorescent-positive single cell clones were plated on individual wells of 96-well plates, pre-coated with 1% (w/v) gelatin to help with cell adherence. Viable clones were expanded, and successful integration of GFP, dTAG or bromoTAG at the target gene locus was confirmed by immunoblotting, PCR and genomic DNA sequencing.

### Immunoblotting

Reduced and denatured cell extracts containing equal amounts of protein (20-30 μg as indicated in legends) were resolved by SDS-PAGE and transferred to nitrocellulose membranes. After blocking membranes in 5% (w/v) non-fat milk in TBS-T (20 mM Tris, 150 mM NaCl, 0.1% Tween-20) for 1 h at room temperature, membranes were incubated with primary antibody diluted in 5% milk TBS-T overnight at 4°C in a shaker. Membranes were then washed 3x10 min in TBS-T with constant shaking and subsequently incubated with HRP-conjugated secondary antibody, also diluted in 5% milk TBS-T solution, for 1 h at room temperature. Membranes were then washed 3x10 min in TBS-T and signal detection was performed via chemiluminescence (Promega) using ChemiDoc imaging system (Bio-Rad).

### Immunoprecipitation

Following determination of protein concentration by BCA protein assay, immunoprecipitation (IP) was performed to pull down proteins of interest. For anti-FLAG IPs, anti-FLAG M2 resin (Sigma-Aldrich) was used; for anti-GFP IPs, anti-GFP Sepharose beads (MRC-PPU Reagents and Services) were used and for Tau IPs, anti-TAU antibody coupled to protein G sepharose beads was used. Pre-IP input and post-IP flowthrough extracts (20 μg protein) were retained to determine IP efficiency. Extracts (0.2-1 mg total protein as indicated) were incubated with 10 μl packed beads (tethered to 10 μg antibodies) overnight at 4°C on a rotating wheel. Beads were collected by centrifugation at 1000 x g for 5 min at 4°C. IPs were subsequently washed 3X with 1 ml lysis buffer and then resuspended in 20 μl of lysis buffer. Pre-IP input, IP and post-IP flowthrough extract samples were reduced in 1X LDS sample buffer (Invitrogen) and boiled at 95°C for 5 min prior to resolving with SDS-PAGE and further analysis.

### In solution digestion of IPs for mass spectrometry analysis

IPs were performed as describe above except after washing the beads three times with 1 ml of lysis buffer, two washes were done with ice-cold PBS. PBS was removed and the beads were incubated with 100 μl of 0.15% trifluoroacetic acid (TFA) for 15 minutes at room temperature (RT) to elute bound proteins. The eluates were transferred to fresh tubes and neutralized using 1 M triethylammonium bicarbonate (TEAB) until the pH was adjusted to approximately 7.0. Subsequently, digestion was performed by adding 1 μg (equivalent) of Promega sequencing-grade trypsin, followed by overnight incubation at 37°C. The next day, digests were acidified and desalted using C18-based StageTips. Peptides were eluted and dried using a vacuum concentrator (Vacufuge) and stored at -80°C until mass spectrometry analysis. The FLAG and GFP IPs were analysed on mass spectrometry using Data Dependant Analysis using the method described below. The FLAG IPs were also analysed using Data Independent Analysis to study PPP2CA interactors.

### In-gel digestion of Tau IPs for mass spectrometry analysis

Anti-Tau IP samples, denatured in 1XLDS sample buffer, were filtered with Corning® Costar® Spin-X® centrifuge tube prior to loading onto SDS-PAGE. The gel was stained with Coomassie InstantBlue® Protein Stain according to manufacturer’s instructions. Gel pieces corresponding to Tau variants (70-130 kDa) were excised and cut into 1x1 mm pieces; gel pieces were destained with 40 mM Triethylammonium bicarbonate and 40% ACN in microcentrifuge tube. Destained gel pieces were dehydrated with 100% ACN for 15 min. Dehydrated gel pieces were subsequently reduced with 5 mM DTT in 40 mM Triethylammonium bicarbonate and incubated at 60°C for 30 min. The reduced gel pieces were alkylated with 20 mM iodoacetamide (IAA) in 40 mM Triethylammonium bicarbonate and incubated for 10 mins in the dark at RT. Alkylated gel pieces were dehydrated with 100% ACN on ice and rehydrated with 10 ng/µl sequencing grade trypsin (Promega) in chilled 40 mM Triethylammonium bicarbonate for 45 min. Once completely rehydrated, excess trypsin was removed, and gel pieces were incubated at 37°C overnight. Peptides were extracted with 100 µl extraction buffer (5% Formic acid; 40% ACN) three times, and 100 µl of 100% ACN was used for final extraction. The peptides were dried completely in Speed-vac concentrator, reconstituted in 30 µl of 0.1% formic acid and desalted on C18 column, before eluting with 30 µl of 40% ACN, 0.1% formic acid twice. Eluted samples were dried completely in Speed-vac concentrator and reconstituted for mass spectrometry analysis.

### Global proteome and phospho-proteome analysis by mass spectrometry

For global total and phospho-proteomic analysis, cells were lysed in urea lysis buffer (8 M urea, 50 mM Triethylammonium bicarbonate buffer (TEAB) pH 8.0, supplemented with protease inhibitors and tablet of PhosSTOP phosphatase inhibitors by sonication with 40% amplitude, on/off 10 second/10 cycles. Extracts were clarified by centrifugation for 20 min at 13,000xg at 4°C. Protein concentration was estimated using the BCA method. Equal protein from each condition was reduced with 5 mM dithiothreitol (DTT) at room temperature for 30 min and alkylated with 20 mM iodoacetamide (IAA) in the dark at room temperature for 15 min. The urea concentration was diluted to less than 2 M with 50 mM Triethylammonium bicarbonate buffer (TEAB) prior to digestion with trypsin (1:20) at 30°C for 16 h. The digestion was stopped with the addition of trifluoroacetic acid (TFA) to give a final concentration of 1% TFA (v/v) and samples were desalted on 200 mg Sep-Pak C_18_ cartridges (Waters). For the Sep-Pak clean-up, the following solvents were prepared fresh: activation solvent (100% (v/v) acetonitrile (ACN)); Solvent-1 (0.1% (v/v) TFA); Solvent-2 (0.1% (v/v) formic acid (FA)); Solvent-3 (50% (v/v) ACN in 0.1% (v/v) FA). Sep-Pak cartridges were equilibrated with 5 mL 100% ACN, followed by 5 mL 50% ACN, 0.1% FA and finally with 5 mL 0.1% TFA twice. Samples were then loaded onto the equilibrated C_18_ cartridges, washed with 5 mL 0.1% TFA four times, followed by washing with 5 mL 0.1% FA. Samples were then eluted with 1 mL 50% ACN, 0.1% FA. Desalted samples were then completely dried in a SpeedVac concentrator. Peptides were dissolved in 100 mM TEAB and labelled using TMT (Thermo Fisher) labels as per the manufacturer’s instructions. Briefly, TMT labels were resuspended in anhydrous acetonitrile, added to assigned samples and incubated for 1 h at room temperature. The labelling reaction was quenched with 5% hydroxylamine for 15 min at room temperature. Labelled peptides from each condition were pooled together and dried. Pooled peptides were resolved by basic reverse phase chromatography fractionation on a C_18_ 250×4.6 mm, 5 μm XBridge column (Waters, Milford, MA) with a flow rate at 500 μL/min with two buffers: buffer A (10 mM ammonium formate, pH 10) and buffer B (80% ACN, 10 mM ammonium formate, pH 10). Peptides were dissolved in 120 μL of buffer A (10 mM ammonium formate, pH10) and separated on a C_18_ reverse phase column by applying a non-linear gradient of 7-40%. A total of 96 fractions were collected and concentrated into 24 fractions for proteomic analysis. 90% sample was used for immobilized metal affinity chromatography (IMAC)-based phospho-peptide enrichment and pooled into 12 fractions. Each concentrated fraction was then dried by SpeedVac. IMAC beads were prepared from Ni-NTA (nitrilotriacetic acid) superflow agarose beads that were stripped of nickel with 100 mM EDTA and incubated in an aqueous solution of 10 mM iron (III) chloride (FeCl_3_) to prepare Fe-IMAC beads. Dried peptide fractions were dissolved to a concentration of 0.5 μg/μL in 80% ACN/0.1% TFA. The mixture of peptides was incubated with beads for 30 min with end-to-end rotation. C_18_ stage tips were prepared by equilibrating with methanol followed by 50% ACN/0.1% FA, and then 1% FA. The beads with enriched peptides were loaded onto C_18_ stage tips and washed with 80% ACN/0.1% TFA. Phosphorylated peptides were eluted from IMAC beads with 500 mM dibasic sodium phosphate, pH 7.0. These peptides were washed with 1% FA before elution using 50% acetonitrile in 0.1% FA. The peptides were then dried by SpeedVac and stored at −80°C until mass spectrometry analysis.

Enriched phospho-peptides and peptides were analysed on an Orbitrap Fusion Tribrid mass spectrometer interfaced with Dionex Ultimate 3000 nanoflow liquid chromatography system. For proteomic analysis, peptides were resolved on an analytical column (Acclaim PepMap RSLC C_18_, 75 μm × 50 cm, 2 μm, 100 Å) at a flow rate of 300 nL/min, using a step gradient of 5-7% solvent B (90% ACN/0.1% FA) for the first 10 min, followed by 7-25% up to 70 min and 25-35% up to 70-85 min. The total run time was set to 100 min. The mass spectrometer was operated in a Data Dependent Acquisition mode in SPS MS3 (FT-IT-HCD-FT-HCD) method. A survey full scan MS (from m/z 400-1400) was acquired in the Orbitrap at a resolution of 120,000 at 200 m/z. The AGC target for MS1 was set as 4 × 10^5^ and ion filling time as 50 ms. The precursor ions for MS2 were isolated using a Quadrupole mass filter at 0.7 Da isolation width, fragmented using a normalized 32% HCD of ion routing multipole and analysed using ion trap. The top 10 MS2 fragment ions in a subsequent scan were isolated and fragmented using HCD at a 65% normalized collision energy and analysed using an Orbitrap mass analyser at a 50,000 resolution, in the scan range of 100-500 m/z. The raw proteomics data were searched using SEQUEST HT search engines with Proteome Discoverer 2.4 (Thermo Fisher Scientific). The following parameters were used for the analysis: Precursor mass tolerance 10 ppm, Fragment mass tolerance 0.1, Enzyme: trypsin, Mis-cleavage: −2, Fixed modification: carbamidomethylation of cysteine residues and TMT of lysine and N-terminal, Dynamic modification: oxidation of methionine. The data were filtered for 1% PSM, peptide and protein level FDR. Only unique peptides were selected for the quantification. Phospho-peptide-enriched fractions from each replicate were searched against the Uniprot protein database using the SEQUEST HT search engines with Proteome Discoverer 2.4 (Thermo Fisher Scientific). A 10 plex TMT reporter ion workflow was loaded, and the following search parameters were used: trypsin protease was selected; two missed cleavages were allowed; oxidation of Met and phosphorylation of Ser/Thr/Tyr were set as variable modifications; and carbamidomethylation of Cys, TMT of lysine and N-terminal were set as fixed modifications. The mass error tolerance for MS1 and MS2 of 10 ppm and 0.02 Da was used. The data were filtered for 1% PSM, peptide and protein level FDR. For identification of the phospho-site probability, the ptmRS node was used. For proteomic data: the SEQUEST output protein group text files were processed using the Perseus software suite^103^. The reporter ion intensities were log2 transformed and the data was normalized by the median for each sample independently. Student-T test was performed and permutation-based false discovery rate of 5% was applied to identify the differentially enriched and significant protein groups. For cluster analysis, the multiple T test ANOVA was carried out with Benjamin Hochberg correction false discovery rate (FDR) of 5%. Significant changes were classified by p < 0.05 and a fold change greater than either 1.5-fold for phospho-proteomic analysis or 2-fold for total proteomic analysis.

### Analysis of FLAG-PPP2CA interactors by mass spectrometry

Following digestion and clean up, the peptides were analysed on an Orbitrap Exploris 480 mass spectrometer interfaced with Dionex Ultimate 3000 nanoflow liquid chromatography system. The dried peptides were reconstituted in 0.1% formic acid and loaded on a trap column (Acclaim PepMap100 C_18_, 100 μm × 2 cm, 5 μm, 100 Å) and separated on an analytical column (Acclaim PepMap RSLC C_18_, 75 μm × 50 cm, 2 μm, 100 Å) at a flow rate of 250 nL/min using a step gradient of 0–8% solvent B (90% ACN/0.1% FA) for the first 12 min, followed by 8–25% up to 110 min, 25-37% for 110-125 min. The total run time was set to 145 min. For DIA MS scan, the resolution was set at 120,000, scan range 375-1500, normalised AGC target to 300%. The ions were fragmented using HCD with stepped energy of 10 and 25, AGC target at 3000%, and variable windows used for MS2 acquisition. The raw files acquired in DIA mode were processed using DIA-NN, version 1.8 (https://github.com/vdemichev/DiaNN). Human Uniprot was used as protein sequence database. Two missed cleavage and a maximum of two variable modifications per peptide were allowed (acetylation of the protein N-termini and oxidation of methionine). Carbamidomethylation of cysteines was set as a fixed modification. This data analysis was carried out using library-free analysis mode in DIA-NN with “deep learning-based spectra and RTs prediction” enabled. MBR was enabled. The resulting data was analysed in Perseus.

### Fluorescence Microscopy and Analysis

Retrovirally transduced SK-N-MC cells were seeded on glass coverslips and subjected to pre-extraction in 0.05% Triton X-100 in PHEM buffer (60 mM PIPES, 25 mM HEPES, 10 mM EGTA, and 4 mM MgSO4·7H20) for 2.5 min at 37°C before fixation in PHEM buffer with 4% PFA for 10 min at 37°C. Fixed cells were then blocked in 5% BSA, 0.1% Triton X-100 in PBS for 1 h at RT followed by incubation with anti-alpha tubulin antibody at a 1:1000 dilution (Invitrogen, MA1-80017) for 1 h at RT. Cells were then washed 3X with PBS and incubated with anti-Rat Alexa FluorTM 647 at 1:1000 (Invitrogen, A78947) for 1 hour at RT. Cells were then washed 3X with PBS and once with Milli-Q water before allowing them to air dry. Coverslips were mounted on glass slides using ProLong TM Glass Antifade Mountant with NucBlue^TM^ (Invitrogen, P36981) and allowed to cure at 4°C overnight. Samples were imaged on a DeltaVison Elite widefield microscope using x100 PlanApoN 1.42 NA objective at 1 × 1 binning and a CoolSNAP HQ camera (Photometrics). Images were deconvolved using Softworks Version 5.5 with enhanced ratio using 10 cycles. Images were then analysed using FIJI. Tubulin signal was used to create a binary mask and colocalized average GFP fluorescence intensity measured. Multiple cells per field of view were analysed with multiple images per condition representative of three independent biological repeats.

### Ternary complex formation - NanoBRET assay

U2OS cells were transiently co-transfected with pBabeD plasmids encoding dTAG and bromoTAG, each tagged at the N- or C-terminus with either NanoLuc or HALO tag, using PEI and seeded onto white 96-well plates. Cells were then treated with either DMSO or BDPIC (200 nM) and HaloTag® 618 Ligand for 24 h before the NanoBRET assay was performed according to the manufacturer’s instructions (Promega, N1661).

### BDPIC competition assay

Tau^bromoTAG/bromoTAG^/^dTAG/+^PPP2CA (Clone 3) cells were seeded onto 6-well plates at 60-70% confluency and incubated overnight. Cells were then treated with either 10 µL DMSO, 250 nM BDPIC, 2500 nM dTAG-V1-neg or 2500 nM cis-AGB1 for 24 h prior to lysis. For competition assay, cells were pre-treated with either 2500 nM dTAG-V1-neg or 2500 nM cis-AGB1 for 1 hour and subsequently treated with 250 nM BDPIC for 24 h. Cells were lysed as described above. Protein concentrations were determined via Bradford assay and cell extracts containing equal amounts of protein (25 µg) were reduced and denatured in 1X LDS with 2% (final) 2-mercaptoethanol for subsequent SDS-PAGE and immunoblotting.

## Supplementary Information

### Supplementary Figures and Legends

**Figure S1:**
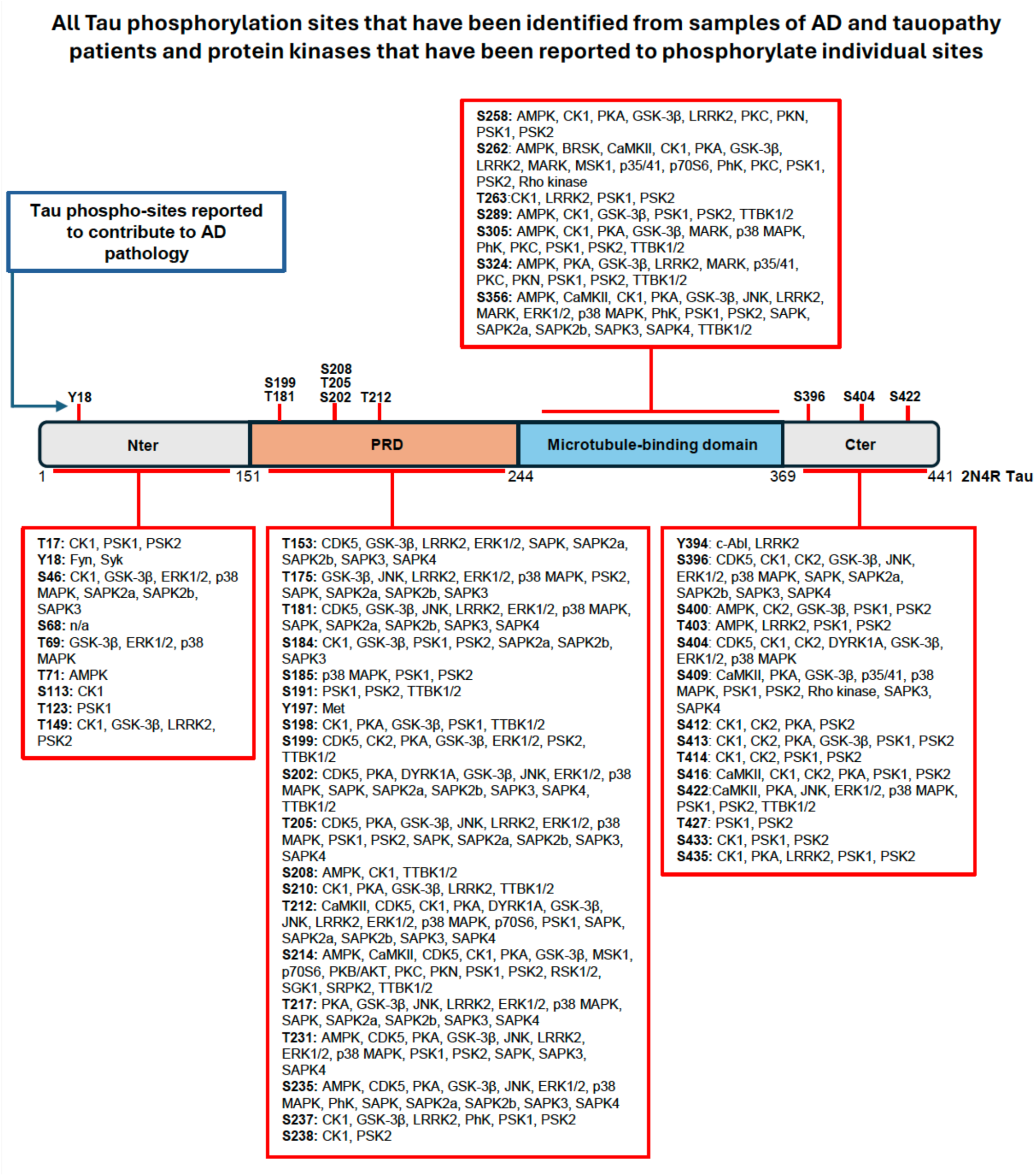
Phospho-atlas of Tau in Alzheimer’s disease (AD). By using a cartoon of 2N4R variant of Tau, all phosphorylated residues on Tau that have so far been reported to have been identified from AD brains are listed. For each phosphorylation site, any kinase that has been previously implicated is also listed. However, it is important to note that for each phospho-site, the stoichiometry of phosphorylation is not known and in different AD patients, the full extent of Tau hyperphosphorylation is impossible to determine. Also denoted above the 2N4R cartoon are the residues whose phosphorylation or phospho-mimetic substitutions have been reported to either enhance the propensity of Tau to aggregate or that these phosphorylation sites have been reported to promote progressive AD pathology. Targeted dephosphorylation of these sites would therefore be potentially beneficial in reversing Tau aggregation and AD progression.

**Figure S2:**
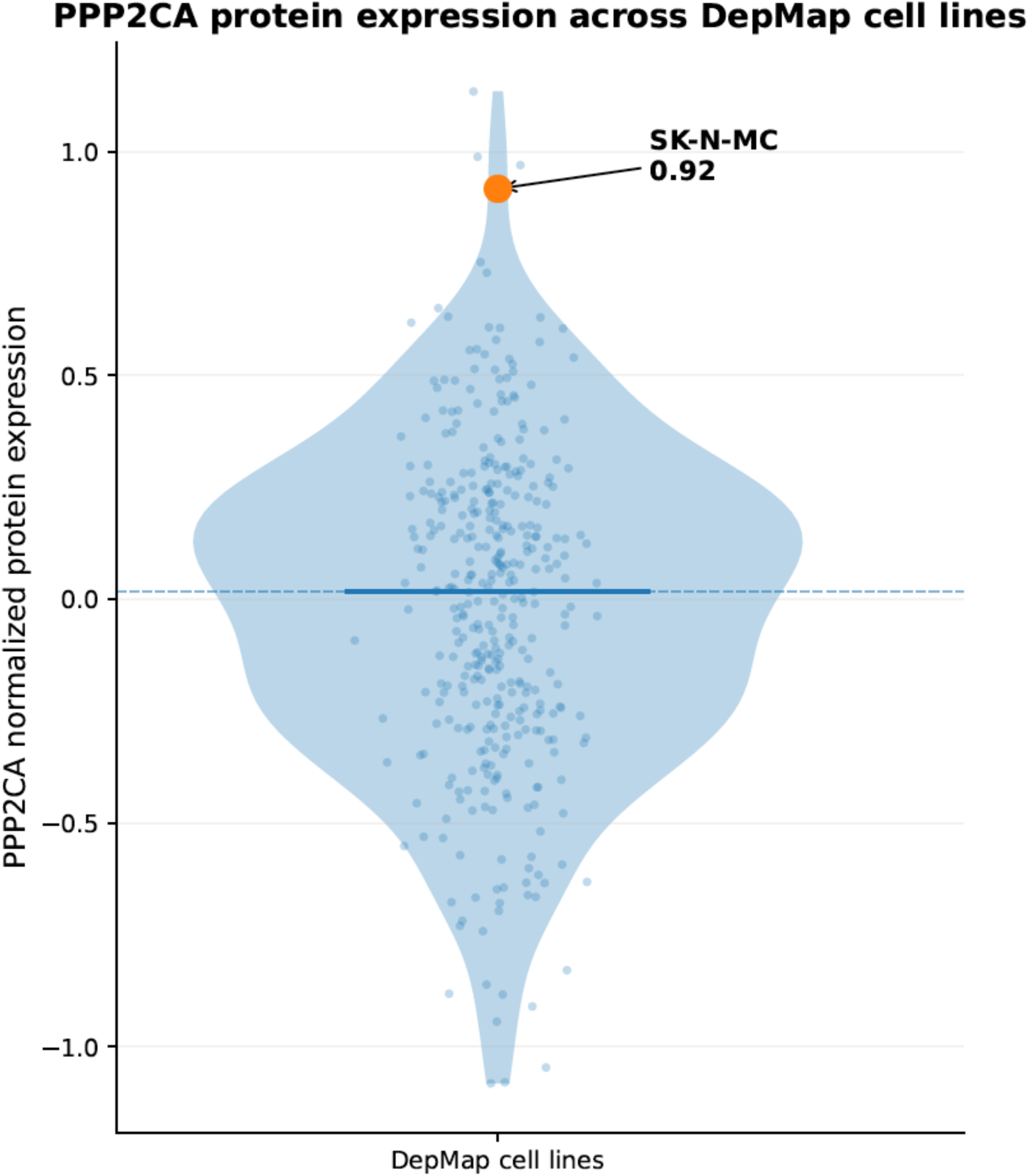
DepMap protein abundance analysis of PPP2CA across 379 cancer cell lines. Cancer Cell Line Encyclopedia (CCLE) proteomics data were mined using DepMap portal and the relative abundance of PPP2CA across all cancer cells (each dot represents a unique cancer cell line) were plotted. SK-N-MC cells were identified as one of the cell lines with highest relative PPP2CA expression.

**Figure S3:**
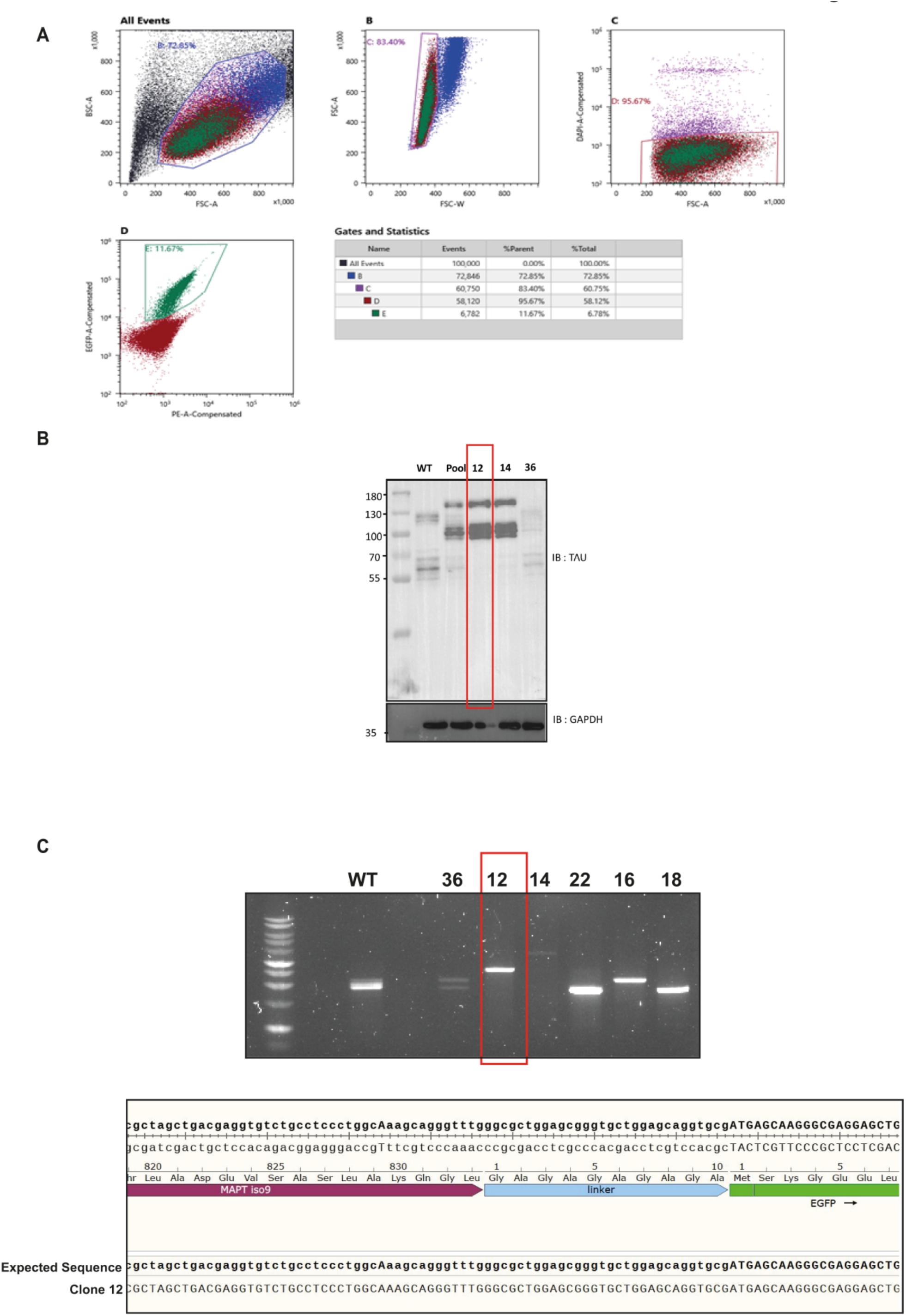
The homozygous insertion of GFP at the native *MAPT* locus by CRISPR/Cas9 genome editing. **(A)** Following two sequential rounds of transfections of SK-N-MC cells with CRISPR/Cas9 genome editing plasmids to insert GFP-tag on *MAPT* locus, GFP-positive cells were sorted by FACS and either pools or single clonal isolates of GFP-positive cells were collected and propagated. **(B)** Wild type SK-N-MC cells, or CRISPR/Cas9 genome edited GFP-positive pool cells, or three individual clonal isolates of GFP-positive cells (clones 12, 14 and 36) were lysed and extracts (20 μg protein) were resolved on 4-12% gradient SDS-PAGE and transferred to nitrocellulose membranes, which were subjected to immunoblotting analysis using the indicated antibodies. Complete upward mobility shift of Tau equivalent to the inserted GFP tag on clones 12 and 14 compared to WT cells indicates a homozygous insertion of GFP on Tau. **(C)** Wild type SK-N-MC cells or clones 36 and 12 were subjected to genomic PCR with primers flanking the C-terminal coding region of *MAPT* and further downstream of the stop codon. A uniform and complete shift of the product on clone 12 compared to WT cells indicated a homozygous insertion of the GFP tag. This in-frame insertion of GFP on Tau was confirmed by DNA sequencing.

**Figure S4:**
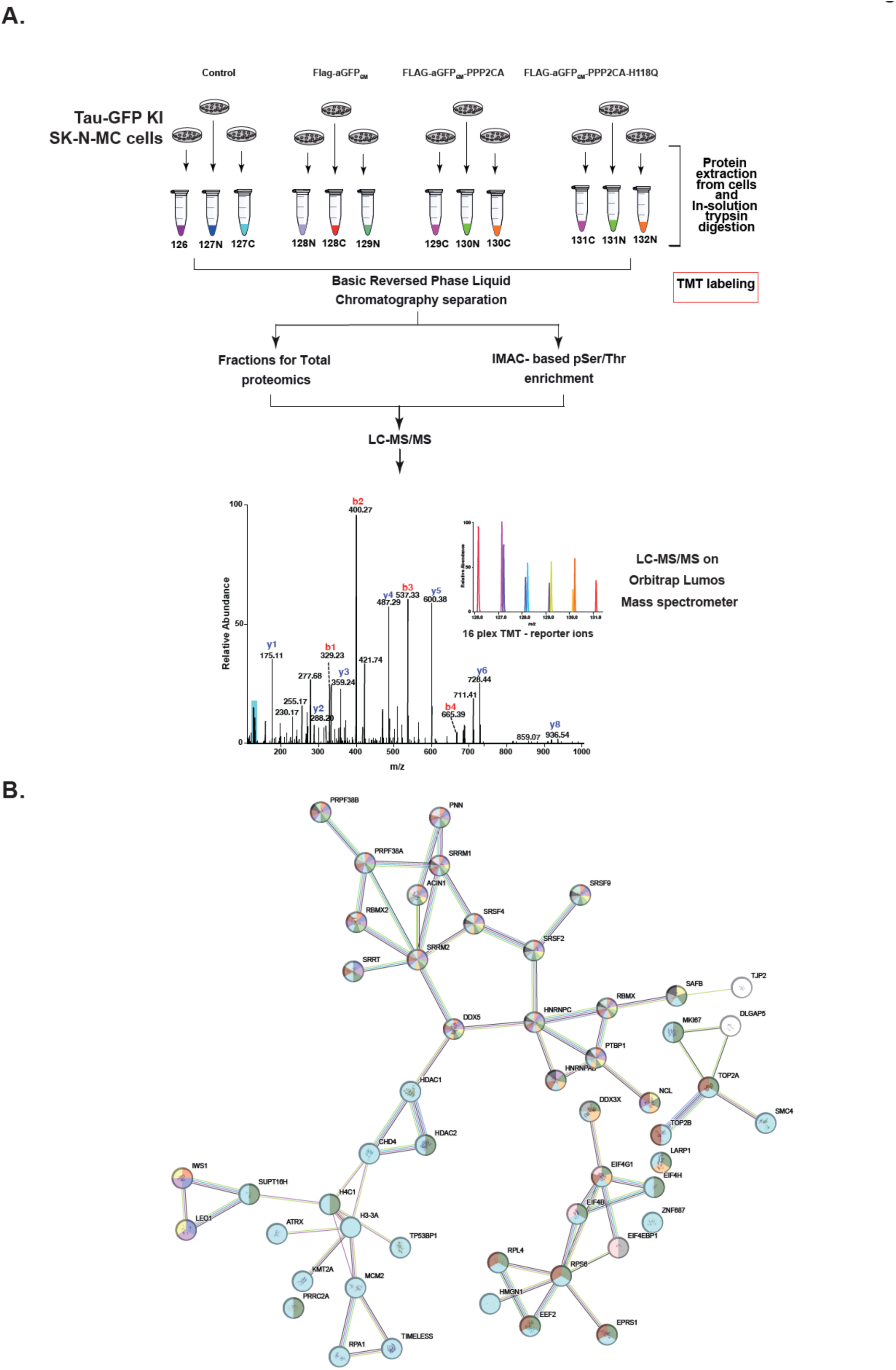
Proteomic workflow to determine the specificity of the AdPhosphatase system. **(A)** Workflow for unbiased quantitative phospho- and total-proteomic analyses in SK-N-MC Tau-GFP KI cells or those stably expressing FLAG-aGFP_6M_, FLAG-aGFP_6M_-PPP2CA or the catalytically inactive FLAG-aGFP_6M_-PPP2CA^H118Q^ mutant. **(B)** String analysis of the proteins that were identified as hypo-phosphorylated when Tau-GFP was targeted for targeted dephosphorylation by FLAG-aGFP_6M_-PPP2CA AdPhosphatase.

**Figure S5:**
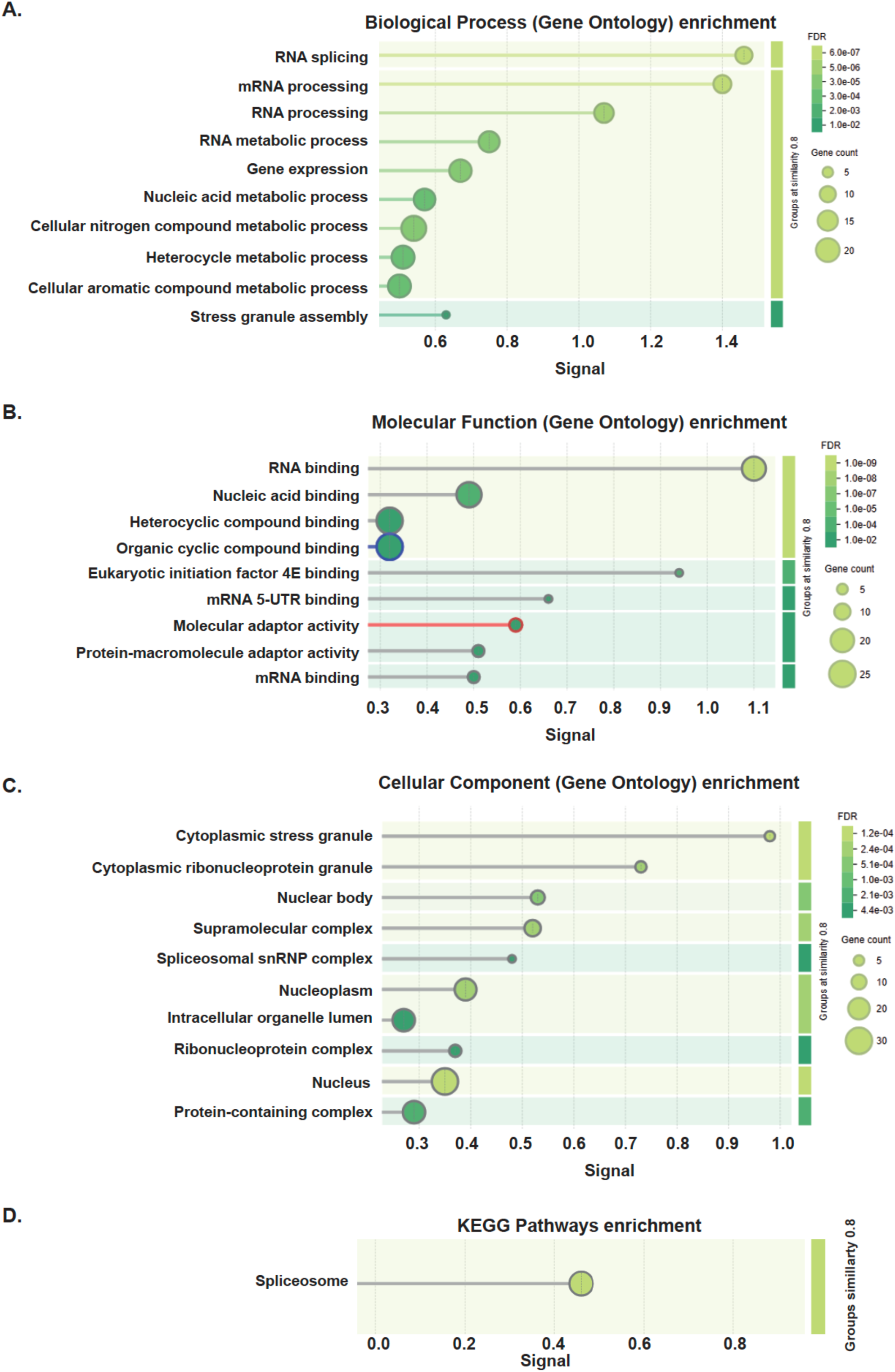
Gene Ontology (GO) enrichment analysis of hypo-phosphorylated proteins when Tau-GFP was targeted for dephosphorylation by FLAG-aGFP_6M_-PPP2CA AdPhosphatase. **(A)** Involvement of hypo-phosphorylated proteins in different biological processes **(B)** Involvement of hypo-phosphorylated proteins in different molecular functions **(C)** Involvement of hypo-phosphorylated proteins in different cellular components **(D)** Involvement of hypo-phosphorylated proteins in KEGG (Kyoto Encyclopedia of Genes and Genomes) pathways.

**Figure S6:**
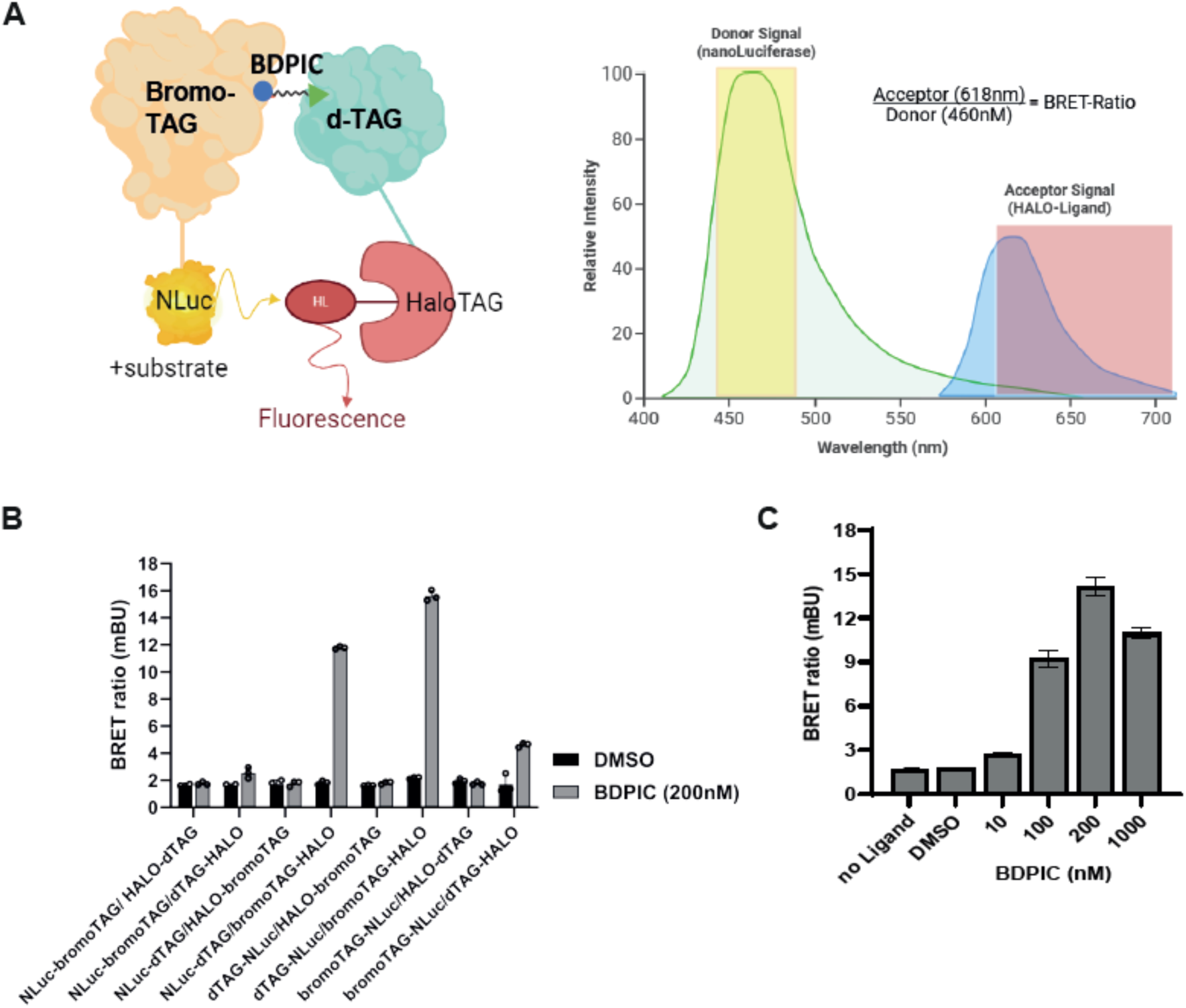
Optimization of NanoBRET Luciferase assay for BDPIC-induced ternary complex formation between bromoTAG and dTAG. **(A)** A cartoon depicting the principles of energy transfer in a NanoBRET luciferase assay. First protein of interest (POI) is expressed as POI1-NanoLuc (NLuc) luciferase fusion. The second POI expressed as POI2-HALO fusion, for which a covalent cell-permeable fluorescent NanoBRET tracer is applied. When POI1 binds POI2 in cells, the tracer also becomes proximal to the NLuc luciferase and provided the proximity is optimal (i.e. a ternary complex has formed), it results in a bioluminescence resonance energy transfer (BRET) signal that can be measured. If BDPIC induces a ternary complex between bromoTAG and dTAG in live cells, using various combinations of NLuc and HALO tags on either protein should allow for determining BRET signals. **(B)** NLuc and HALO tags were attached to bromoTAG and dTAG in either N- or the C-termini in all combinations and expressed in cells. Luciferase and BRET signals were measured following treatment of cells with DMSO or BDPIC and BRET ratios plotted for each pair of constructs. Clear differences in BRET ratios induced by BDPIC compared to DMSO treatment are evident for NLuc-dTAG:bromoTAG-HALO, dTAG-NLuc:bromoTAG-HALO and bromoTAG-NLUC:dTAG-HALO combinations. **(C)** dTAG-NLuc and bromoTAG-HALO expressing cells were subjected to treatment with DMSO or the indicated doses of BDPIC as indicated for 24 h prior to lysis and BRET measurement, which are plotted.

**Figure S7:**
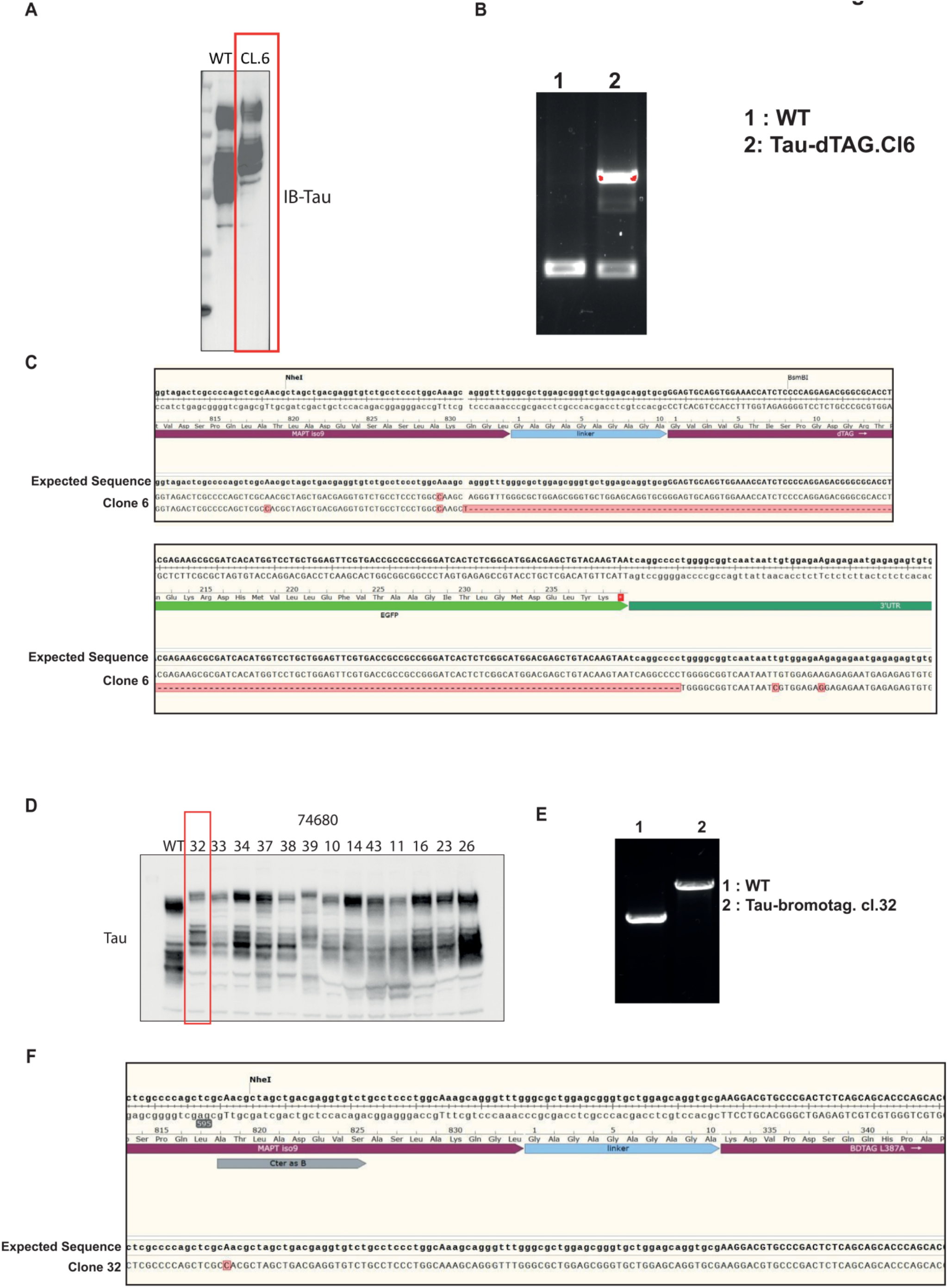
Validation of Tau-dTAG and Tau-bromoTAG KI in SK-N-MC cells by CRISPR/Cas9 genome editing. **(A)** Following two sequential rounds of transfections of SK-N-MC cells with CRISPR/Cas9 genome editing plasmids to insert in-frame dTAG-IRES-GFP cassette on *MAPT* locus, GFP-positive cells were sorted by FACS. Of multiple GFP-positive clones isolated (not shown), Clone 6 (boxed) shows a electrophoretic mobility shift equivalent to the molecular weight of dTAG on all Tau variants compared to WT SK-N-MC cells, as shown by Tau immunoblotting. **(B&C)** Analysis of WT and Tau-dTAG KI (Cl6) SK-N-MC cells by PCR (B) and genome sequencing (C) show that dTAG was knocked in on one *MAPT* allele, while the second allele was modified also modified but in such a way that no native molecular weight Tau protein expression was observed. This Tau-dTAG KI (Cl6) SK-N-MC cell line was used for further studies. **(D)** As in (A), except a donor plasmid to insert in-frame bromoTAG-IRES-mCherry cassette on *MAPT* locus was used and 13 mCherry-positive single cell clones collected, propagated and analysed by immunoblotting. Homozygous Tau-bromoTAG knock-in was confirmed in several clones by immunoblotting with anti-Tau antibody, which showed a clear upward mobility shift equivalent to the molecular weight of bromoTAG compared to Tau from wild type (WT) SK-N-MC cells. **(E&F)** WT or Tau-bromoTAG KI (Cl32) SK-N-MC cells from (D) were subjected to PCR (E) or genomic sequencing (F) analyses, which confirmed a homozygous insertion of bromoTAG on both *MAPT* alleles. This clone (Cl32) has been used for all subsequent studies.

**Figure S8:**
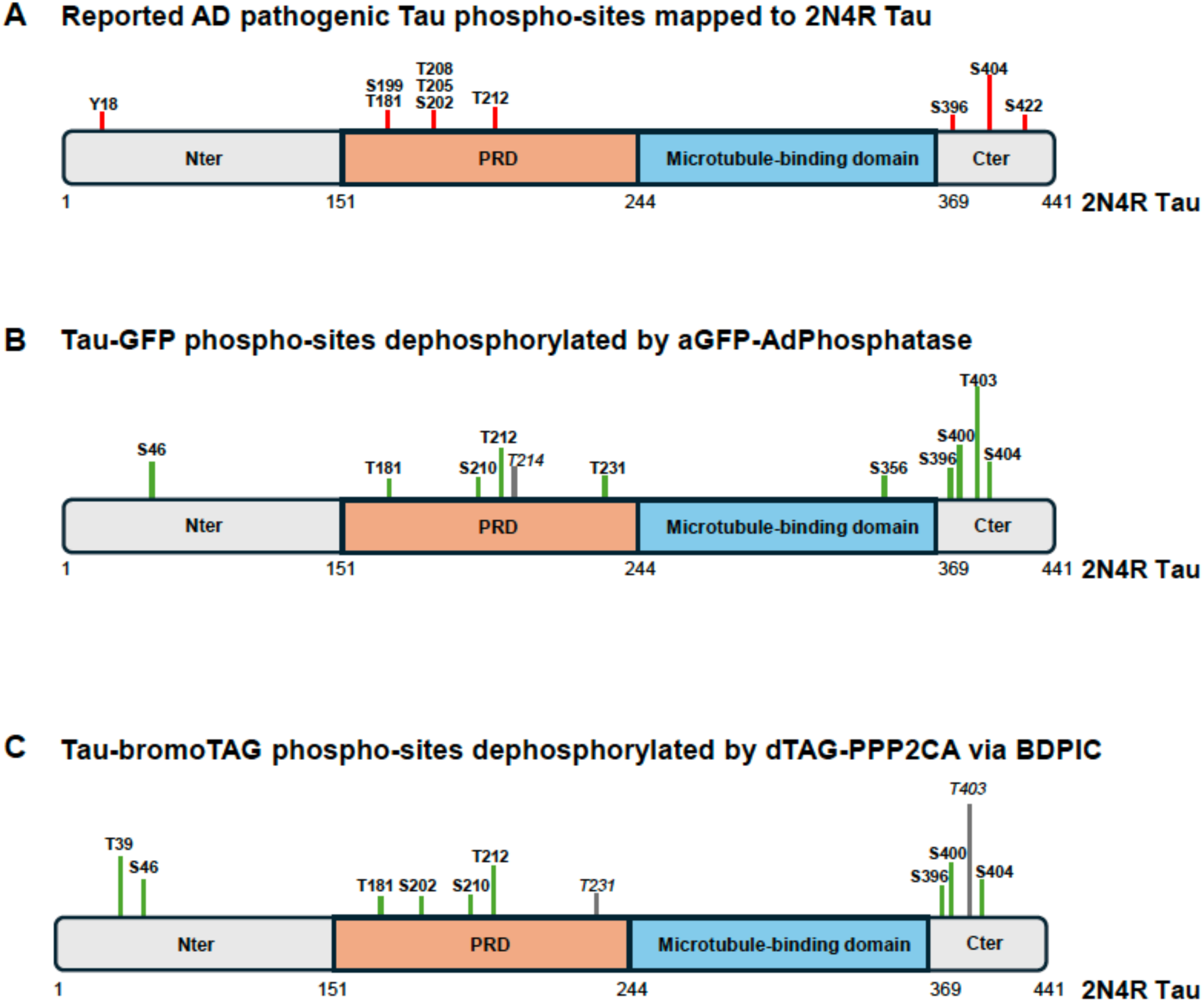
Phospho-sites mapped on 2N4R variant of Tau. **(A)** The reported pathogenic phospho-sites on Tau: This panel provides an annotated linear map of the human 2N4R Tau protein highlighting the major pathogenic phosphorylation sites reported to be associated with Tauopathies and Alzheimer’s disease. **(B)** Tau-GFP phospho-sites targeted for dephosphorylation by the aGFP-AdPhosphatase indicated on 2N4R Tau. **(C)** Tau-bromoTAG phospho-sites targeted for dephosphorylation by dTAG-PPP2CA via BDPIC indicated on 2N4R Tau.

**Figure S9:**
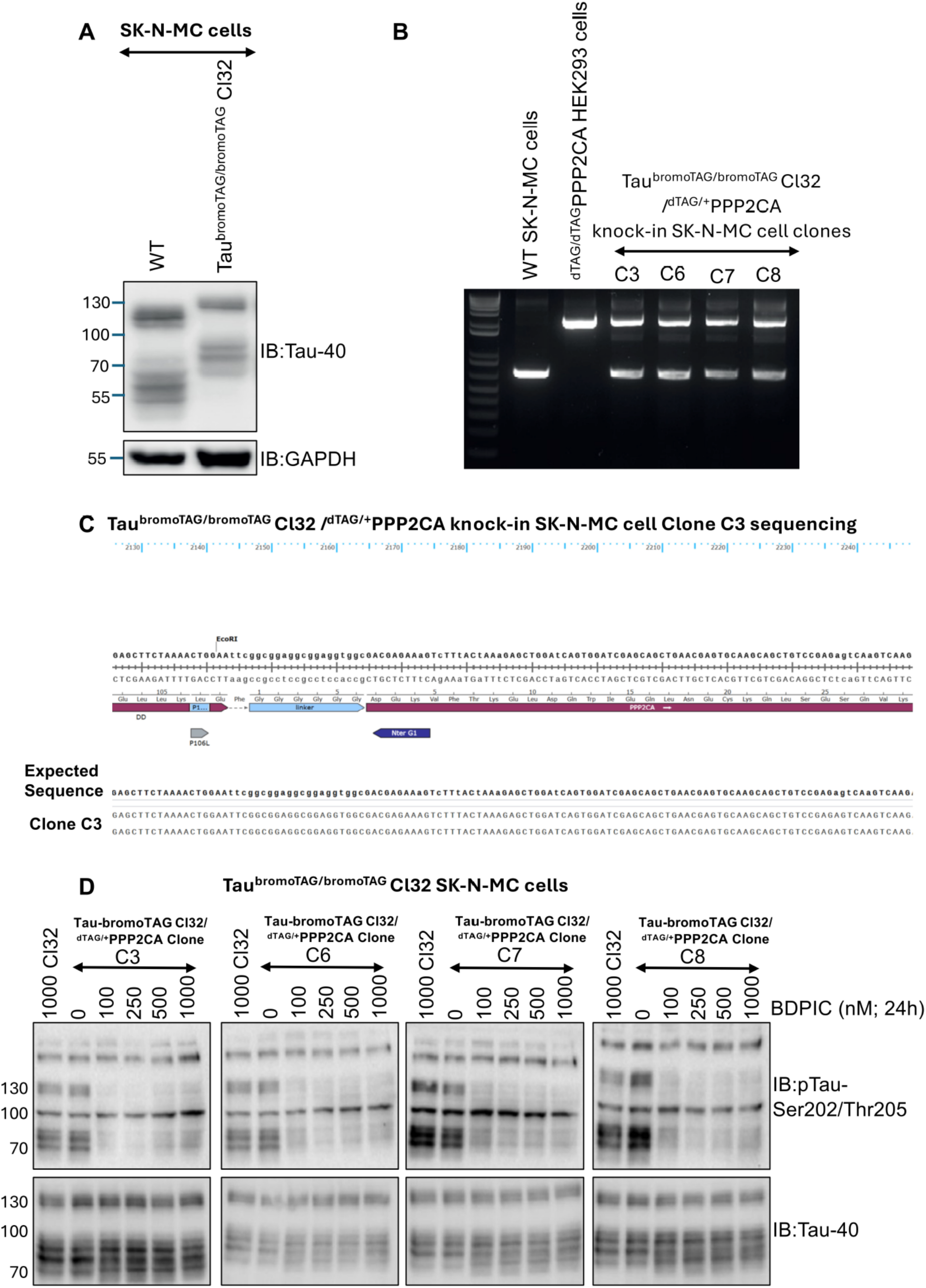
Heterozygous insertion of dTAG on *PPP2CA* locus in *Tau^bromoTAG/bromoTAG^* KI (Clone 32) SK-N-MC cells by CRISPR/Cas9 genome editing. **(A)** WT or Tau^bromoTAG/bromoTAG^ KI (Cl32) SK-N-MC cells were lysed and extracts (20 μg) subjected to SDS-PAGE and transfer to nitrocellulose membranes, which were immunoblotted with the indicated antibodies. Homozygous Tau-bromoTAG knock-in is demonstrated by a clear upward mobility shift of Tau equivalent to the molecular weight of bromoTAG compared to Tau from WT SK-N-MC cells. Tau^bromoTAG/bromoTAG^ (Cl32) was used to knock in dTAG on *PPP2CA* locus. **(B)** *Tau^bromoTAG/bromoTAG^* knock-in (Cl32) SK-N-MC cells were subjected to CRISPR/Cas9 genome editing to insert dTAG *(FKBP12^F36V^)* on *PPP2CA* locus using a GFP-IRES-dTAG donor cassette. The indicated GFP-positive clones were subjected to genomic DNA isolation and PCR to assess successful insertion. Genomic DNA derived from homozygous *^dTAG/dTAG^PPP2CA* knock-in HEK293 cells was employed as positive control for PCR. PCR showed heterozygous dTAG-PPP2CA KI in all clones, as signal corresponding to WT allele was still visible in all clones. **(C)** The heterozygous dTAG knock-in on *PPP2CA* locus was confirmed on Clone C3 by DNA sequencing. **(D)** Indicated *Tau^bromoTAG/bromoTAG^/^+/dTAG^PPP2CA* double KI SK-N-MC cell clones were treated with increasing concentration of BDPIC (0-1000 nM) for 24 h prior to lysis. As control, parental *Tau^bromoTAG/bromoTAG^* (Cl32) SK-N-MC cells were treated with 1000 nM BDPIC for 24 h prior to lysis. Extracts (25 μg protein) were resolved on 4-12% gradient SDS-PAGE and transferred to PVDF membranes, which were subjected to immunoblotting analysis using the indicated antibodies.

